# Human antigen-specific memory natural killer cell responses develop against HIV-1 and influenza virus and are dependent on MHC-E restriction

**DOI:** 10.1101/2020.11.09.374348

**Authors:** Stephanie Jost, Olivier Lucar, Taylor Yoder, Kyle Kroll, Sho Sugawara, Scott Smith, Rhianna Jones, George Tweet, Alexandra Werner, Phillip J. Tomezsko, Haley L. Dugan, Joshua Ghofrani, Marcus Altfeld, Adam Grundhoff, Michaela Muller-Trutwin, Paul Goepfert, R. Keith Reeves

**Author notes:** Corresponding author R. Keith Reeves, Center for Virology and Vaccine Research, Beth Israel Deaconess Medical Center, 3 Blackfan Circle, Boston, MA 02215. Current address: Committee on Immunology, University of Chicago, Chicago, IL 60637, USA.

## Abstract

For over a decade, multiple studies have disputed the notion of natural killer (NK) cells as purely innate lymphocytes by demonstrating that they are capable of putative antigen-specific immunological memory against multiple infectious agents including two critical global health priorities – HIV and influenza. However, the mechanisms underlying antigen specificity remain unknown. Herein, we demonstrate that antigen-specific human NK cell memory develops upon exposure to both HIV and influenza, unified by a conserved and epitope-specific targetable mechanism firmly dependent on the activating CD94/NKG2C receptor and its ligand HLA-E, and confirm these findings by three rigorous and novel assays. We validated the permanent acquisition of antigen-specificity by individual memory NK cells by single-cell cloning. We identified biomarkers of antigen-specific NK cell memory through RNA-Seq transcriptomic fingerprints and complex immunophenotyping by 30-parameter flow cytometry showing elevated expression of KLRG1, α4β7 and NKG2C. Finally, we show individual HLA-E-restricted peptides that may constitute the dominant response in HIV-1- and influenza-infected persons in vivo. Our findings clarify the mechanisms behind formation of antigen-specific memory NK cells, and suggest they could be targeted for future vaccines, cure strategies, or other therapeutic interventions.

## INTRODUCTION

Natural killer (NK) cells are lymphocytes classically considered part of the innate immune system based on their ability to mediate very rapid and nonspecific responses to virally infected or neoplastic cells. A plethora of studies have provided compelling evidence for the significant contribution of NK cells to the immune control of HIV and influenza virus infections^1–3^, two of the most critical global health priorities. NK cells represent a potent antiviral effector cell population that can promptly respond to HIV and influenza exposure without the need for prior antigen sensitization. The interest in harnessing NK cell functions for vaccine design and therapeutic interventions against these pathogens has dramatically increased in recent years, largely driven by a series of observations indicating subpopulations of NK cells, called memory or adaptive NK cells, manifest multiple different forms of durable adaptive capabilities^4–20^. This includes reports of true antigen-specific memory NK cells^5,7,8,10,13,14,18–20^ that can mediate recall responses against multiple infectious agents including HIV^7,13,18^ and influenza^7,14^. Adaptive NK cell subsets have been associated with protective effects in people living with HIV (PLWH)^21–24^ and exposure to influenza antigens induces protective influenza-specific memory NK cells in mouse models^7,14^. Thus, antigen-specific memory NK cells represent a third arm of the immune system that can be targeted by prophylactic or therapeutic interventions.

To date, only a few studies support the existence of true antigen-specific recall responses by NK cells in humans, which have been described against cytomegalovirus (CMV), varicella zoster virus (VZV), hepatitis A (HAV) and B (HBV) virus and bacillus Calmette-Guerin^15,18–20,25,26^ and the opacity surrounding the mechanisms of memory NK cell formation represents a major obstacle to harnessing adaptive NK cell functions. Herein, we provide proof that antigen-specific NK cell memory develops upon exposure to HIV and influenza in humans, and also provide the first mechanistic evidence for memory NK cell recognition of target antigens at the single-cell and single-peptide level.

## RESULTS

### Evidence for human NK cells mediating HIV- and influenza antigen-specific responses

Clear evidence of HIV- and influenza-specific recall responses by NK cells in humans is lacking. We first tested HIV antigen specificity using a modified intracellular cytokine staining (ICS) assay designed to uniquely quantify CD107a+ and IFN-γ+ HIV Gag-specific peripheral blood NK cells in viremic PLWH or healthy donors (Fig. 1a, Table S1). In PLWH, 0.5 to 6% of NK cells were reactive against HIV Gag peptides, mirroring responses found in SIV-infected rhesus macaques^13^, whereas in healthy donors, NK cell responses were generally undetectable above background. The capacity of NK cells isolated from PLWH to specifically react against HIV antigens was further confirmed using a flow cytometry-based cytotoxicity assay. To do so, we assessed the killing of autologous B lymphoblastoid cell lines (BLCL) pulsed with a pool of HIV Gag peptides by untouched NK cells isolated either from PLWH or from healthy donors (Fig. 1b, Fig. S1). Significant anti-HIV Gag activity by NK cells could only be detected in PLWH, while killing of BLCL pulsed with a pool of EBV, CMV and influenza peptides was comparable between PLWH and healthy individuals. Potent lysis of MHC-devoid K562 cells, a target cell line devoid of MHC commonly used to evaluate NK cell cytotoxicity and that triggers non antigen-specific stimulation, was similarly achieved by NK cells from PLWH and HIV-negative donors. PLWH included patients on combination antiretroviral therapy (cART) with undetectable viral loads, untreated viremic PLWH, as well as HIV elite controllers who achieve spontaneous control of viral replication in the absence of treatment (HIV RNA levels <50 copies/mL for at least a year) (Table S1)^27^. NK cells from elite controllers with detectable HIV gag-specific cytotoxic activity displayed the most potent responses observed. These data showed that NK cells that can specifically respond to HIV antigens develop in PLWH and mediate potent anti-HIV activity, suggesting HIV-specific NK cells might significantly contribute to control of HIV. Using a similar ICS-based assay as described above, we next measured the ability of human NK cells to mediate antigen-specific responses to influenza and CMV, to which most humans have pre-existing immunity memory. Enriched NK cells from donors were stimulated with peptide pools derived from the conserved matrix protein 1 (MP1) and nucleoprotein (NP) of the 2009 H1N1 pandemic strain (A/California/08/2009 and A/California/04/2009, respectively) or derived from CMV pp65 (Fig. 1c). We detected influenza positive NK cell responses in 45% of HIV-negative donors who provided blood samples between 2017 and 2020. Altogether, these results suggested that peripheral blood NK cells can mediate antigen-specific recall responses in humans, a hallmark of memory.

**Fig. 1.**
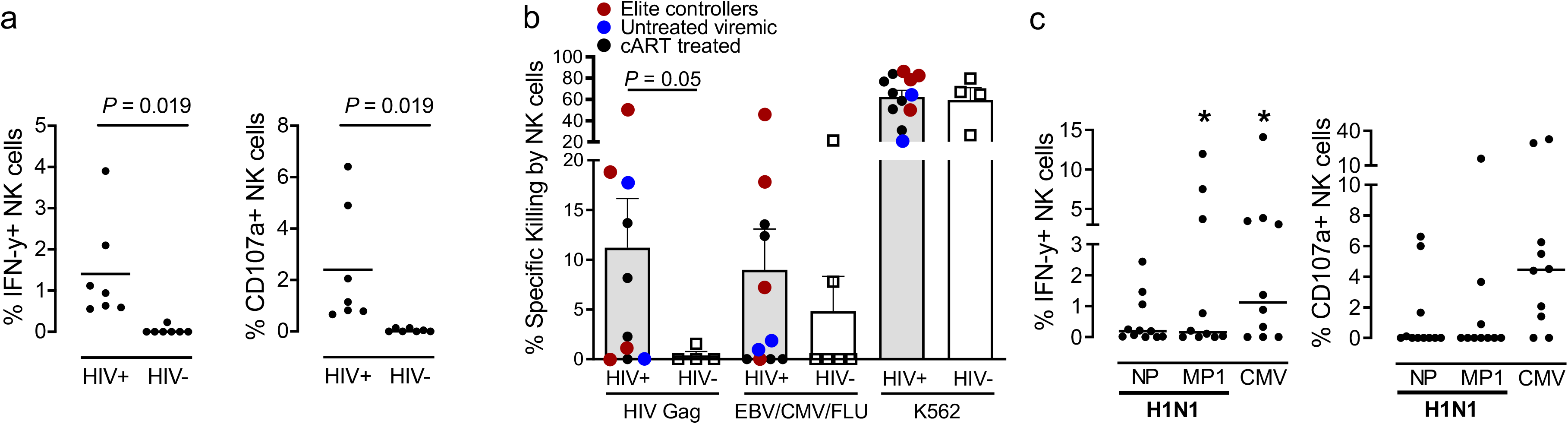
Human NK cells mediate antigen-specific responses against HIV and influenza. **a,** Enriched NK cells from PLWH or healthy donors were co-cultured with autologous BLCL that had been pulsed with 2ug/mL peptide pools derived from HIV Gag (HIV-1 Consensus B; provided by the NIH AIDS Reagent Program) and NK cell responses assessed by ICS. Dead cells were excluded. Dot plots show proportions of IFN-y+ and CD107a+ NK cells after subtracting background (unstimulated). Asterisks indicate significant differences between healthy donors and samples from PLWH. **b,** Autologous BLCL were pulsed with a pool of HIV Gag overlapping peptides or with the CEF (CMV, EBV and influenza) control peptide pool and were labeled with the CellTrace Violet dye. Mock-pulsed BLCL serving as intra-well controls were labeled with the green dye CFSE. Purified NK cells were co-cultured with BLCL at 5:1 E:T ratios (equal mixture of pulsed target BLCL and non-pulsed control BLCL) for 16 hours, and specific lysis of BLCL was determined by flow cytometry. Killing of HLA-deficient K562 cells was used as additional positive control. **c,** Enriched NK cells 11 HIV-negative healthy donors were incubated overnight with 2ug/mL peptide pools derived from influenza A/California/04/2009(H1N1) NP and A/California/08/2009(H1N1) MP1 or CMV pp65 and NK cell responses assessed by ICS. Dead cells were excluded. Dot plots show proportions of IFN-y+ and CD107a+ NK cells after subtracting background (unstimulated). Asterisks, significant differences compared to unstimulated controls. * p<0.05.

### Single-cell NK cloning confirms antigen-specificity of memory NK cells

To further investigate the existence of antigen-specific memory NK cells that can efficiently respond to HIV and influenza and study these likely rare cells in more detail, we clonally expanded individual peripheral blood NK cells from 20 PLWH (4 cART-treated and 16 viremic untreated) and 8 healthy adults (Fig. S2). Using a cytotoxicity assay, we assessed the ability of single NK cell clones (NKCL) to lyse MHC-devoid K562 cells (to confirm normal NK cell function) or autologous BLCL pulsed with a pool of self-peptides, CMV pp65-, HIV Gag-, HIV Envelope (Env)-, A/California/04/2009 H1N1 NP- and A/California/08/2009 H1N1 MP1-derived overlapping peptides (Fig. 2). As expected, NKCL could potently lyse K562 cells and had little to no reactivity to BLCL pulsed with a self-peptide control. Approximately 29% of NKCL from PLWH (Fig. 2a) and 45% of NKCL from healthy adults (Fig. 2b) displayed positive responses against HIV or influenza, respectively. While the magnitude of HIV-specific responses did not differ between treated and untreated PLWH (Median % Gag-specific killing, cART-treated: 6, viremic: 4.8, p=0.43; median % Env-specific killing, cART-treated: 10, viremic: 7.4; p=0.35), HIV-specific NKCL were more frequently detected in untreated viremic participants (% HIV-specific NKCL, cART-treated: 20, viremic: 32). Strikingly, we were able to isolate NKCL with very robust anti-HIV (up to 87% specific lysis against Env) and anti-influenza H1N1-(up to 57% specific lysis against NP) cytotoxic activity. Interestingly, 30% of NKCL displaying over 6% specific killing against HIV Gag or Env were able to respond to both antigens, with NKCL presenting the highest responses (over 30% specific killing) being able to respond to either Gag or Env but never both. Together these data provide the first proof of HIV- and influenza-specific NK cells in humans and show that NK cells displaying true and robust antigen-specific killing are found at the single cell level.

**Fig. 2.**
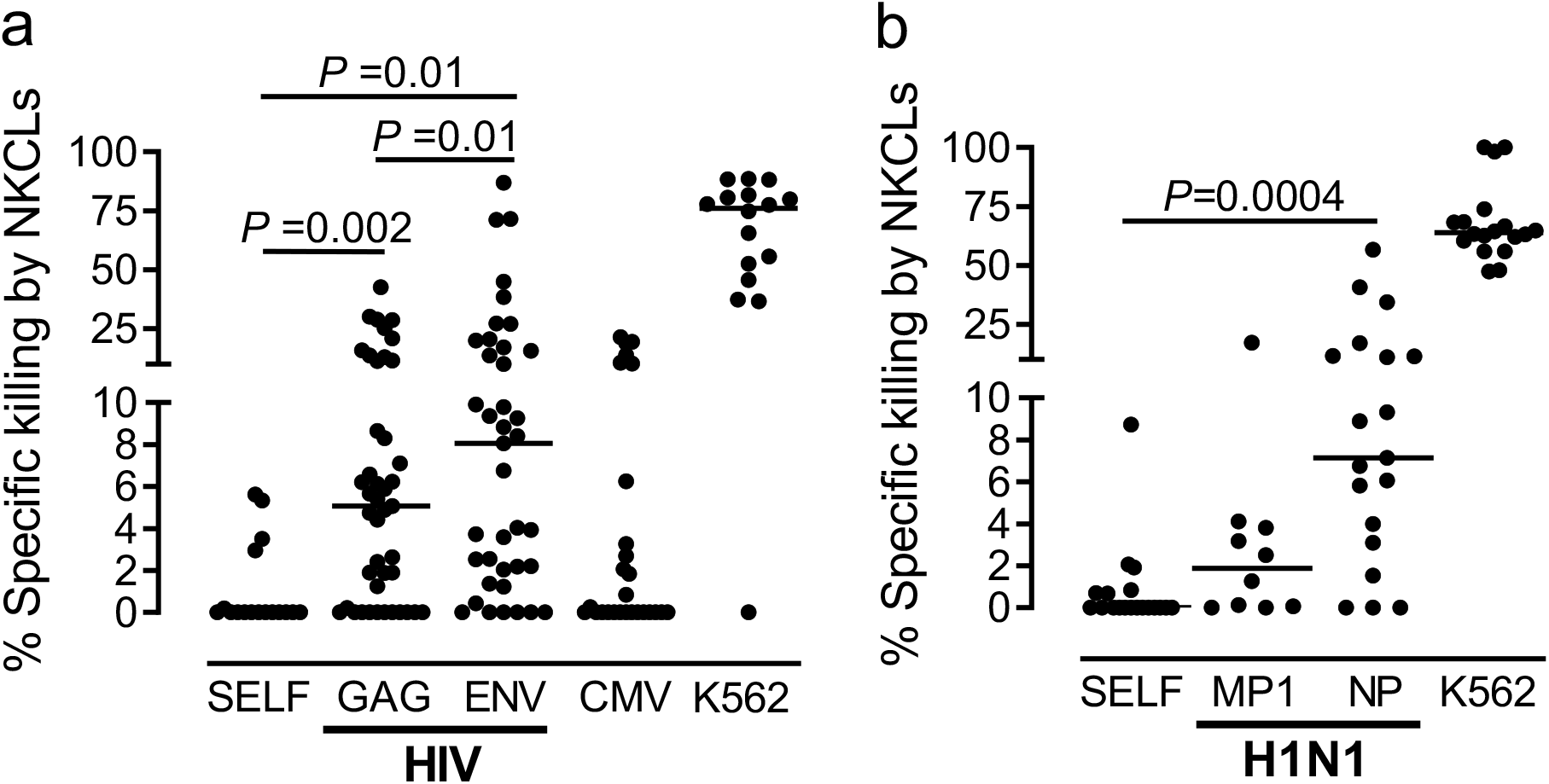
Single-cell cloning of human antigen-specific NK cells. Percentages of antigen-specific lysis by **a,** 46 (out of 159) NKCL from 20 PLWH that reacted against HIV and **b,** 19 (out of 40) NKCL from 8 healthy donors that reacted against influenza. CAM cytotoxicity assays were used to evaluate lysis after co-culture of NKCL with autologous BLCL pulsed with indicated peptide pools. Non-specific lysis was assessed by measuring killing of mock-pulsed autologous BLCL. NKCL reactive against HIV antigens were identified as NKCL mediating specific killing at least twice above specific killing of self-peptides-pulsed B-LCL after background subtraction, and, for those with no self-peptide available data, at least twice above non-specific killing (mock pulsed B-LCL). NKCL found unreactive or over-reactive based on those criteria are not displayed.

### Mechanisms of memory NK cell target recognition and killing are dependent on the activating NKG2C receptor

To define transcriptional profiles associated with antigen-specific NK cell responses, we first performed RNAseq analysis on 8 NKCL with anti-HIV activity and 7 NKCL that did not react against HIV from the same individuals. Overall, 12 genes were significantly upregulated and 4 genes downregulated in HIV Env/Gag-specific NKCL (Fig. 3a, Fig. S3). Of interest, HIV-specific NKCL expressed *NLRP7* and *GSTM2*, which are involved in cellular defense responses, as well as genes involved in regulation of the actin cytoskeleton, such as *FRMD5, CDC42EP4, PLEKHG2*, and in metabolic functions, such as *RGPD8* and *Ak5*. Pathway enrichment analysis supported clear changes in metabolic pathways in HIV-specific NKCL. Overall, our analysis revealed transcriptional profiles consistent with maturation and activation that are associated with memory NK cells endowed with potent anti-HIV activity, but also identified a transcriptional profile that may be unique to this cell type.

**Fig. 3.**
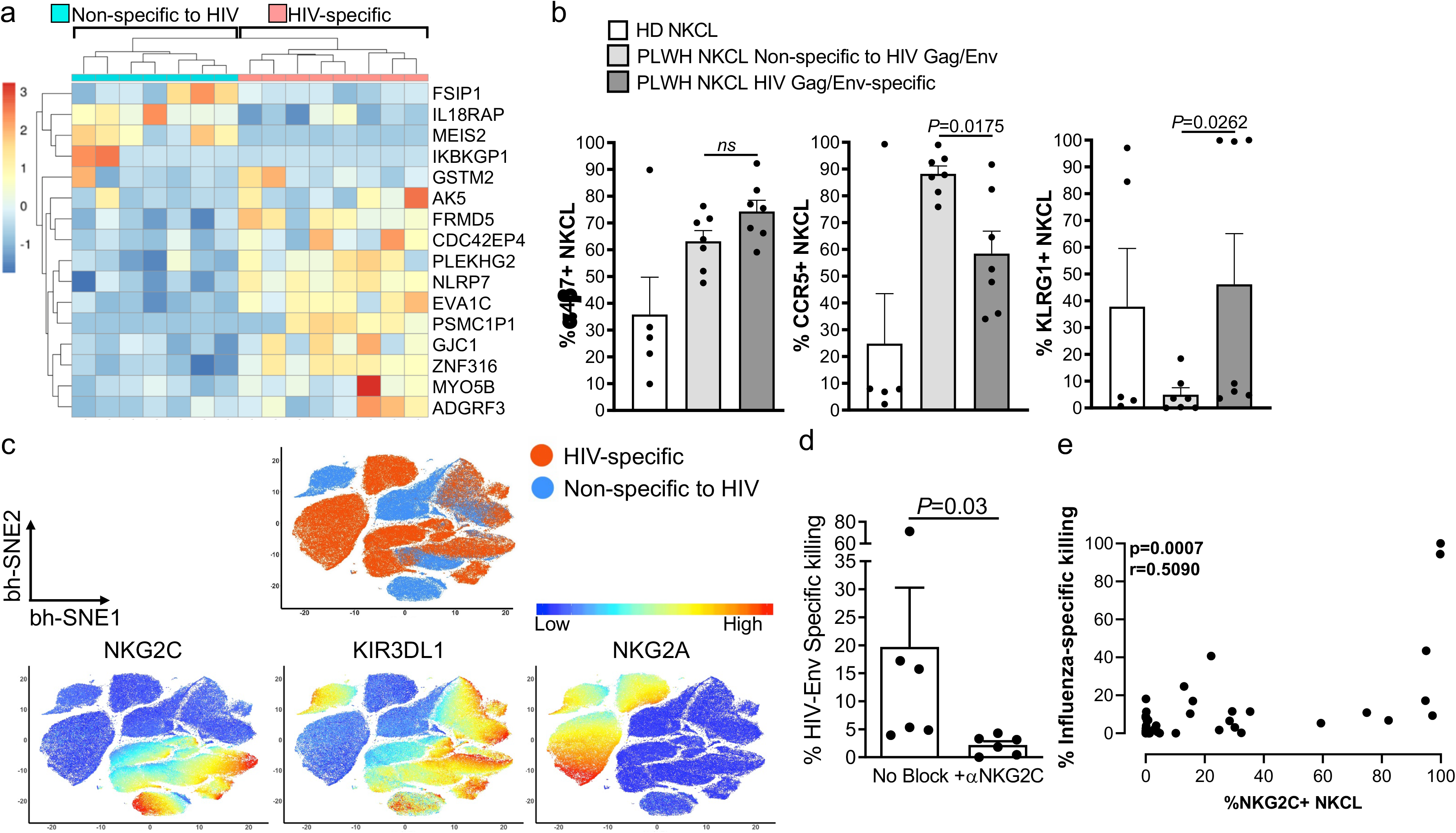
Human antigen-specific NK cell responses are associated with NKG2C expression. **a,** Hierarchical clustering and heatmap of the 16 differentially expressed genes distinguishing HIV-specific NKCL (n=8) and NKCL not reacting to HIV Gag/Env (n=7) from the same untreated viremic PLWH. Normalized Z-scores are displayed as a colorimetric scale of blue (lower expression) to red (higher expression). **b,** Bar graphs show mean + SEM proportions of NKCL expressing α4β7, CCR5 and KLRG1 evaluated by flow cytometry and comparing 5 NKCL from 2 healthy donors (HD) with 14 NKCL from one untreated viremic PLWH, including 7 HIV Gag/Env-specific NKCL and 7 NKCL that did not react to HIV Gag/Env. **c,** Non-linear t-SNE (bh-SNE) plots of 14 NKCL from one untreated viremic PLWH, showing distinct clustering for 7 HIV reactive and 7 non HIV-reactive NKCL, with HIV-reactive NKCL being partly associated with cell clusters expressing high levels of the inhibitory NKG2A or high levels of activating NKG2C and/or inhibitory KIR3DL1. **d,** Antigen-specific killing of HIV Env-pulsed BLCL by 6 NKCL in the presence of isotype control or NKG2C blocking antibodies. Bars represent mean+SEM. **e,** Spearman correlation analysis between frequencies of NKG2C+ NKCL and specific killing of BLCL pulsed with influenza NP, MP1 or HA.

To further define the subset of peripheral blood NK cells that drives antigen-specific memory responses, we then compared expression levels of 21 cell surface receptors between NKCL with anti-HIV Env/Gag activity and those that did not react to those HIV antigens using advanced multiparameter flow cytometry (Table S2). Phenotypic analysis revealed that most HIV-specific NKCL express activating (i.e, NKp46, NKG2D, NKG2C) and inhibitory (NKG2A, CD85j) receptors typically found on NK cells, trafficking markers (i.e., α4β7, CXCR6, CCR5, CCR7), co-stimulatory molecules (i.e., 2B4, CD2, CD8) and various levels of maturation/activation markers (CD57, KLRG1, PD1, Tim-3, HLA-DR) or KIRs (Fig. S4). Compared to NKCL that did not react to HIV, HIV-specific NKCL had enhanced expression of KLRG1 and α4β7 and lower expression of CCR5 (Fig. 3b). Furthermore, analysis using non-linear dimensionality reduction algorithm (*t*-SNE) revealed distinct clustering for HIV reactive and non-HIV-reactive NKCL, with HIV-reactive NKCL being partly associated with cell clusters expressing high levels of the inhibitory NKG2A or high levels of activating NKG2C and/or inhibitory KIR3DL1 (Fig. 3c and Fig. S5).

NKG2C is an activating NK cell receptor that forms heterodimers with CD94 and interacts with the nonclassical MHC class I molecule HLA-E bound to HLA-E-stabilizing peptides. High cell surface expression of NKG2C/CD94 has been consistently linked to long-lived NK cells endowed with adaptive capabilities that develop upon human CMV infection^11,28–31^. Variants of CMV UL-40-derived peptides that mimic canonical HLA class-I-derived leader peptides are presented by HLA-E and finely tune adaptive NKG2C+ NK cell functions^15,26^, suggesting HCMV-specific recognition via the NKG2C/HLA-E axis. Moreover, we previously showed that in non-human primates, HIV antigen-specific memory NK cell responses largely depend on NKG2A/C^13^. As HIV-specific NKCL were associated with high NKG2C expression (Fig. 3c), we directly assessed the role of NKG2C in HIV antigen-specific memory NK cell responses. HIV Env-specific NKCL were co-cultured with BLCL pulsed with HIV Env peptides in the presence of blocking antibodies against activating receptors and specific lysis measured (Fig. 3d, Fig. S6a). NKG2C blockade significantly decreased HIV Env-specific responses by NKCL, while blockade of natural cytotoxicity receptors did not or only marginally impacted HIV-specific NK cell responses, corroborating our previous findings^13^. Accordingly, we found a positive correlation between influenza-specific killing by NKCLs and the proportions of NKG2C+ NKCL (p=0.0007, r=0.5090) and NKCL clones displaying over 10% specific killing against influenza antigen had significantly higher NKG2C expression than NKCLs that did not or only marginally reacted against influenza (Fig. 3e, Fig. S6b). Altogether, our results confirm that antigen-specific memory NK cell responses are dependent on NKG2C, likely indicative of an HLA-E-dependent recognition mechanism.

### HIV- and influenza-derived peptides bind HLA-E, the ligand for NKG2C, and potently activate antigen-specific NK cells

HLA-E displays limited polymorphism and presents largely conserved nonameric peptides derived from leader sequences of other HLA class I molecules. However, pathogen-derived peptides^32–35^, including HIV Pol, Vif and p24^36–38^ have been shown to bind HLA-E. To establish the relevance of an HLA-E-dependent recognition mechanism in antigen-specific memory NK cell responses against two unrelated viruses, we first screened for canonical peptides derived from HIV Env, HIV Gag and influenza NP that can stabilize HLA-E at the surface of K562 cells stably expressing HLA-E*0101 (Fig. 4a, Fig. S7, Table S3). Among all tested nonameric peptides (84 for HIV Env, 60 for HIV Gag, 23 for influenza NP), two HIV Env-derived and two influenza NP-derived peptides were identified that could stabilize HLA-E either to the same extent as the positive control (CMV UL40 VMAPRTLIL) or up to two-fold greater.

**Fig. 4.**
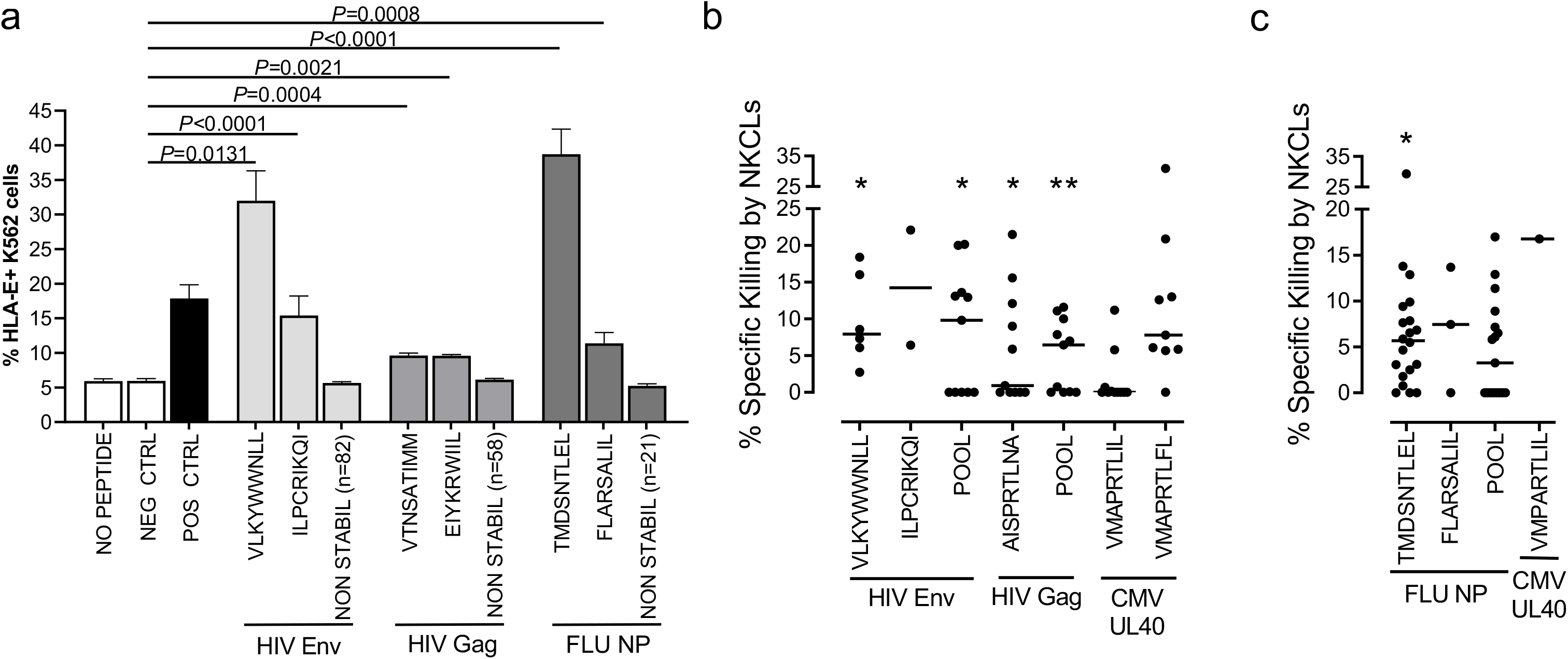
Human antigen-specific memory NK cell responses are mediated via recognition of HLA-E-binding viral peptides. **a,** K562 cells stably expressing HLA-E*0101 were either left mock pulsed or pulsed for 16h with 10ug/mL of nonameric peptides derived from HIV consensus B Gag (60 total) or Env (84 total) or from A/California/04/2009(H1N1) NP (23 total). Controls included CMV pp65-derived NLVPMVATV that do not stabilize HLA-E (NEG), and VMAPRTLIL, a CMV/HLA-Cw3 leader sequence-derived peptide that stabilizes HLA-E (POS). HLA-E surface stabilization was assessed by flow cytometry. Bars represent mean percentages of HLA-E+ K562 cells + SEM for indicated individual peptides or from all non-stabilizing peptides (n) pooled from at least 3 distinct experiments. Dot plots represent percentages of antigen- or peptide-specific lysis by 17 NKCL from 5 untreated viremic PLWH **(b)** and 20 NKCL from 5 healthy donors **(c)** after subtracting non-specific lysis (killing of mock pulsed autologous BLCL). CAM cytotoxicity assays were used to evaluate lysis after co-culture of NKCL with autologous BLCL pulsed with peptide pools encompassing the whole HIV Env, HIV Gag or H1N1 NP sequence or with indicated single HLA-E-binding nonameric peptides derived from CMV UL40, HIV Gag, HIV Env, or H1N1 NP. Asterisks, significant differences compared to mock-pulsed pulsed prior to background subtraction. * p<0.05, ** p<0.01.

To evaluate the effect of HLA-E-binding peptides on antigen-specific NK cell function, we compared killing by NKCL from 5 PLWH of BLCL pulsed with pools of 15-mer peptides with 11 amino acids overlap covering the whole HIV Gag and HIV Env with that of BLCL pulsed with single HLA-E-binding peptides, including HIV Env-derived VLKYWWNLL and ILPCRIKQI or HIV Gag AISPRTLNA, a peptide that has been previously shown to significantly bind HLA-E and affect NK cell function^36,37^ (Fig 4b). 4 out of 11 NKCL (36%) and 5 out of 6 NKCL (83%) reacted to single HLA-E-binding HIV Gag- and HIV Env-derived peptides (range of positive responses twice above the background: 6-21% Gag-specific and 6-22% Env-specific killing). Similarly, we evaluated killing of BLCL pulsed with either pools of 15-mer overlapping peptides encompassing the whole NP sequence or single TMDSNTLEL and FLARSALIL peptides (all derived from A/California/7/2009 H1N1), by 20 NKCL from 5 healthy donors (Fig. 4c). Thirteen NKCL (65%) reacted to TMDSNTLEL, among which 6 also displayed responses against the NP pool, two of these being also able to respond to FLARSALIL (range of positive responses twice above the background: 3-29% NP-specific killing). These results show that HIV Env- and influenza NP-derived nonamers can bind HLA-E and potently activate patient-derived memory NKCL.

Next, we sought to confirm the *in vivo* relevance of the novel peptide responses we identified among single-cell clones. First, we showed stimulation with newly identified HIV Env- and influenza NP-derived HLA-E-binding peptides alone promotes CD107a upregulation on a subset of primary NK cells in PLWH and healthy donors, respectively (Fig. 5). CD107a up-regulation in NK cells correlated significantly with NK cell-mediated cytotoxicity^39,40^ and in NHPs, we also found a correlation between HIV-specific killing by memory NK cells and CD107a up-regulation by memory NK cells. Thus, our results strongly indicate *in vivo* activation of cytotoxic antigen-specific adaptive NK cells by single HLA-E binding nonamers. Collectively, our work provides the first recognition mechanism underlying antigen-specific recall responses mediated by NK cells in humans, which have the potential to be harnessed for vaccine design or other therapeutic interventions.

**Fig. 5.**
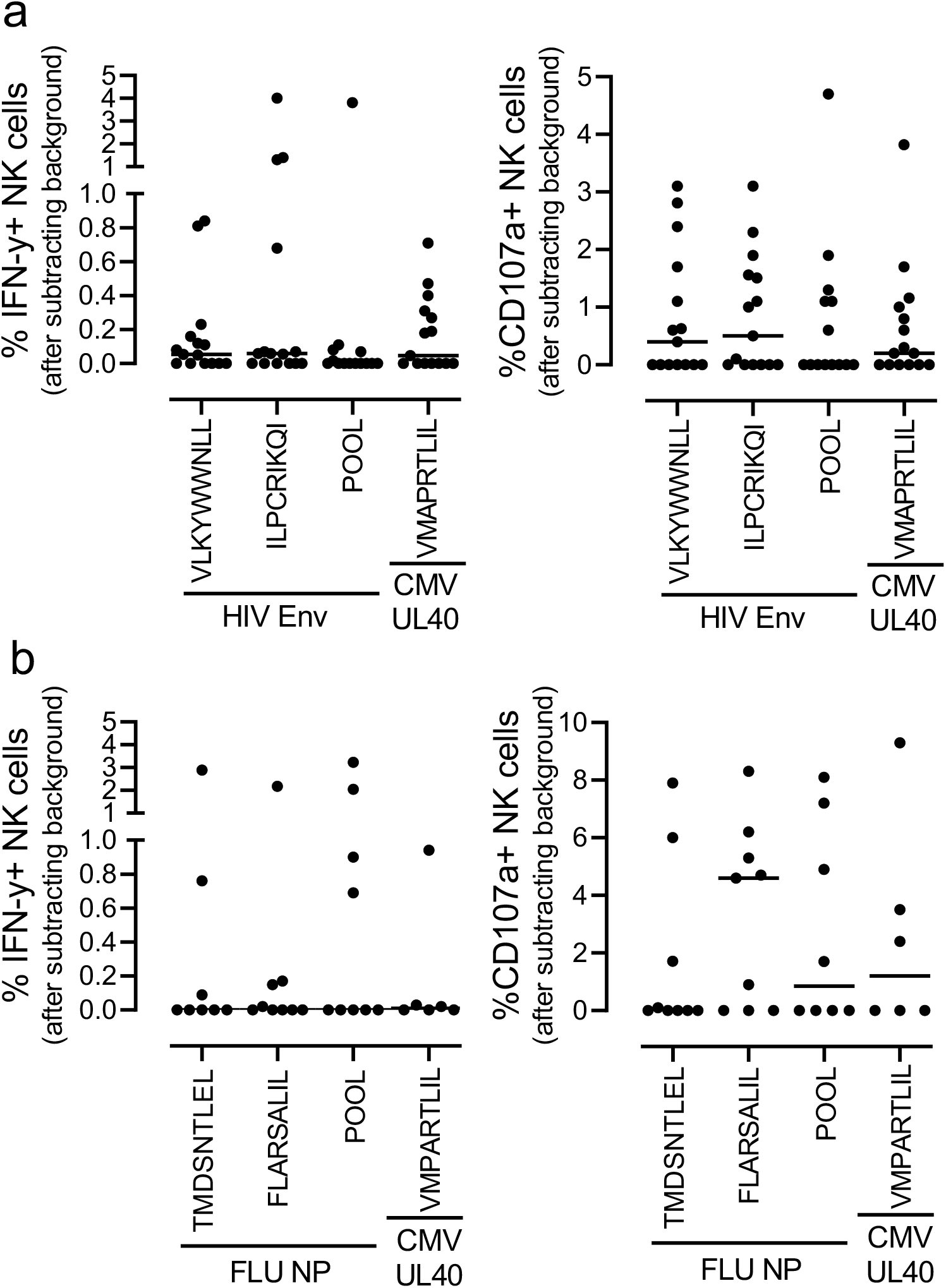
HIV Env- and H1N1 NP-derived HLA-E-binding peptides elicit primary NK cell responses. **a,** PBMCs samples from 15 PLWH (11 cART-treated and 4 untreated viremic) were incubated for 16h with 2μg/mL of 1 pool of 15 amino acid peptides overlapping by 11 amino acids and spanning HIV Env or single HIV Env- and CMV UL40-derived nonamers in the presence of CD107a antibodies. GolgiStop and GolgiPlug were added for the last 2h incubation prior to surface and intracellular cytokine staining. **b**, Purified NK cells from 9 healthy donors were incubated for 4h with autologous BLCL pulsed with 2μg/mL of 1 pool of 15 amino acid peptides overlapping by 11 amino acids and spanning H1N1 NP or single H1N1 NP- and CMV UL40-derived nonamers in the presence of CD107a antibodies, GolgiStop and GolgiPlug. Dot plots represent proportions of IFN-y producing (left panel) or actively degranulating, cytotoxic NK cells, as judged by cell surface expression of CD107a (right panel) after subtracting proportions of IFN-y+ and CD107a+ unstimulated NK cells, respectively. Bars represent the median.

## DISCUSSION

Thus far, antigen-specific memory responses mediated by NK cells in humans have been described in VZV-experienced adults^18^ as well as in HBV-infected or HAV/HBV-vaccinated individuals^19,20^. While the idea of primate NK cell memory has become well-accepted among immunologists, until now there have been no reported mechanisms describing how NK cells recognize and distinguish individual antigens. Our study provides the first mechanistic evidence of human NK cell memory against pathogens of global public health importance that cannot currently be efficiently prevented by vaccination. We show that HLA-E possesses the previously unappreciated ability to bind HIV and influenza peptides and activate virus-specific NK cells. HLA-E has gained significant attention in the vaccine field as broad *Mamu*-E restricted CD8^+^ T cell responses have been implicated as immune correlates of protection in macaques vaccinated with a cytomegalovirus-vectored SIV vaccined^41,42^, and this overall function may be conserved between rhesus macaque *Mamu*-E and human HLA-E^43^. Vaccine strategies that mobilize HLA-E could induce HIV- or influenza-specific memory NK cells that recognize broad epitopes across viral antigens. One major issue for the development of an HIV vaccine or a universal influenza vaccine is the considerable variability of viral strains circulating among humans. Antigen-specific NK cell responses relying on activation of the NKG2C/HLA-E axis is in accordance with the fact that in our studies, some antigen-specific NKCL were able to potently react to several antigens from the same as well as from different viruses. Further, we present primary HLA-E-restricted responses for both pathogens which may represent dominant memory NK cell responses in vivo that could be further targeted. This feature could be highly beneficial for vaccine design and allow a limited number of memory NK cells to target a broader range of HIV and influenza strains than memory T cells.

In addition to the lack of mechanism previously described for NK cell memory, the field also continues to search for comprehensive biomarkers to distinguish adaptive subpopulations of NK cells from classical NK cells. CD49a, CXCR6, NKG2C, and suppressed gamma signaling chain have also been implicated to partially delineate these populations, but particularly for true antigen-specific memory NK cells these phenotypes maybe be incomplete^44^. Herein, we describe multiple phenotypes that advance biomarker usage for NK cell memory. KLRG1 has been described as a marker of NK cell activation^45^, which defines subsets of memory-like NK cells with protective functions against Mycobacterium tuberculosis^46^ and subsets of NK cells that mediate antigen-specific memory responses against HBV antigens^19^. Upregulation of KLRG1 on HIV-specific NKCL further suggest KLRG1 might be a useful biomarker to identify antigen-specific memory NK cells among bulk NK cells. Additionally, α4β7 was found to be upregulated on HIV-specific NKCL and this corroborates previous observations that HIV and SIV infection specifically mobilizes mucosal-homing NK cells^47,48^. Not surprisingly, due to our mechanism described in these studies and others, NKG2C was also highly expressed on memory NK cells. However, antigen-specific memory NK cell responses likely do not solely depend on the NKG2C/HLA-E axis, and other pathways may be additive or alternative to this mechanism. In particular, activating KIRs represent another important family of receptors that may regulate memory NK cell function. Specific KIR genes expressed in conjunction with their HLA ligands have been clearly associated with improved HIV, HCV, CMV, flavivirus, and bacterial infection control^49–52^, and while the interaction of inhibitory KIRs with HLA class I ligands has been studied in detail, ligands for most activating KIRs remain elusive. Besides triggering NK cell activation through interactions between HLA class I ligands and activating NK cell receptors, influenza- or HIV-derived peptides may also disrupt inhibitory signals mediated through NKG2A or other inhibitory molecules and HLA-E or their cognate HLA class I ligand, respectively^37,53–57^, further promoting NK cell responses. This possible mechanism is consistent with HIV-reactive NKCL being partly associated with high expression levels of the inhibitory NKG2A and KIR3DL1 receptors (Fig. 3c and Fig. S5). Future studies beyond the scope of this work will be required to determine if KIR mediated recognition is complementary or independent to this highly dominant HLA-E dependent mechanism.

Finally, various mechanisms have been proposed to explain the spontaneous ability to maintain undetectable viral loads in the absence of cART in HIV elite controllers. Notably, specific variations in the human HLA class I gene locus, highly functional CD8 T cell and NK cell responses and distinct configuration of the proviral reservoir have been associated with enhanced virological control^58–61^. Our data now suggest enhanced HIV-specific NK cell cytotoxicity as a potential correlate of viral control in HIV elite controllers and further investigations are warranted to better define the role played by antigen-specific NK cell responses in this rare subset of PLWH.

Overall, we addressed two major deficits to the field: we demonstrate that a conserved and epitope-specific targetable mechanism dependent on NKG2C/HLA-E underlies true antigen-specific memory NK cell responses in humans, and together with previous observations, our detailed phenotypic and transcriptional analyses suggest complex phenotypes may be the most effective at tracking antigen-specific memory NK cells *in vivo* rather than individual markers.

## MATERIALS AND METHODS

### Generation of functional autologous BLCL

EBV-transformed B cell lines (BLCL) were generated as autologous targets for antigen-specific NK cell assays using standard protocols. Briefly, PBMC for each donor were incubated with EBV supernatant 37°C for 1 hour. After washing, PBMC were cultured for 3-6 weeks in R10 supplemented with 1 ug/ml CSA, being split approximately every 7 days. After achieving log phase growth, for which maximum antigen processing and presentation occurs, BLCL were used in subsequent assays. BLCL functional capacity was confirmed by processing of DQ-Ovalbumin DQ assay (Molecular Probes). Briefly, BLCL were incubated with 1mg/mL DQ for 2 hours. DQ-O cleavage was confirmed by flow cytometry analysis on an LSRII instrument against negative controls including BLCL without DQ-O and with DQ-O added but kept at 4 degrees.

### Flow cytometry-based antigen-specific NK cell killing assay

K562 cells and autologous BLCL were labeled with 1uM CellTrace™ Violet Cell Proliferation Kit (Invitrogen) for 20 minutes according to manufacturers’ instructions. CellTrace Violet-labelled BLCL were then pulsed for 1.5 hours at 37C with 2ug/mL HIV Gag (HIV-1 Consensus B; by the NIH AIDS Reagent Program) or CEF (CMV, EBV and influenza)-derived overlapping peptide pools in culture medium consisting in RPMI-1640 supplemented with 2 mM L-glutamine, 100 ug/ml streptomycin, 100 U/ml penicillin and 5% human serum, washed with PBS and resuspended in culture medium. Non-pulsed BLCL served as internal control and were labeled with 2uM CellTrace™ CFSE Cell Proliferation Kit (Invitrogen) for 15 minutes according to manufacturers’ instructions, washed and resuspended in culture medium. NK cells were purified using AutoMACS NK cell enrichment kit and resuspended in culture medium with 0.1ng/mL recombinant human IL-15. 350,000-400,000 NK cells were co-cultured with BLCL at 10:1 E:T ratio containing an equal mixture of pulsed target cells and non-pulsed control cells or K562 cells for 16 hours at 1M/mL, and subsequently stained using CD3-A700 (UCHT1), CD19-BV711 (HIB19), CD16-BV785 (3G8) and CD56-PEcy7 (B159). Specific lysis of BLCL was calculated as follows: (% sample lysis with NK effectors - % basal lysis without NK effectors) / (100 - % basal lysis without NK effectors) (Fig. S2).

### Analysis of primary NK cell or NKCL responses to HIV and influenza peptide antigens by intracellular cytokine staining

To measure primary antigen-specific NK cell responses, cryopreserved PBMCs from healthy donors or PLWH were thawed and used either unfractionated, after CD3+ T cell depletion using the EasySep Human CD3 Positive Selection Kit II and the and the protocol for Using EasySep Positive Selection Kits for Cell Depletion provided by the manufacturer (STEMCELL Technologies) or after NK cell enrichment using the EasySep Human NK Cell Enrichment Kit (STEMCELL Technologies). To measure antigen-specific NKCL responses, NKCL were never cryopreserved and used directly from cultures after phenotypic analysis to confirm expression of CD16, CD56 in the absence of CD3. Autologous BLCL were pulsed for 1.5 to 2 hours at 37C with 2ug/mL of peptide pools consisting of 15-mer sequences with 11 aa overlap covering the complete sequence of the influenza virus A/California/04/09(H1N1) NP (PepTivator, Miltenyi), A/California/08/2009(H1N1) MP1 (PepMix, JPT Peptide Technologies), CMV pp65 (NIH AIDS Reagent Program), HIV Env and HIV Gag (both from NIH AIDS Reagent Program) or with 5ug/mL of single nonameric peptides derived from influenza NP, HIV Env, HIV Gag or CMV UL40 (Thermofisher). After washing, peptide-loaded BLCL were co-cultured with enriched NK cells or NKCL at a 1:1 E:T ratio at 37C for 4h to 15h with CD107a BV786 or CD107a FITC (H4A3) (BD Biosciences). In some instances 1ng/mL of recombinant human IL-15 (R&D) was also added. Unfractionated PBMC from PLWH or CD3-depleted PBMCs from healthy donors were stimulated directly with peptides without addition of BLCL. 1uL/mL GolgiPlug (BD Biosciences) and 0.7uL/mL GolgiStop (BD Biosciences) were added for the whole 4h of incubation or, when incubation lasted over 4h, for the last 2h of incubation. Cells were stained first with the LIVE/DEAD^®^ Fixable Blue Dead Cell Stain Kit (Invitrogen), then with CD3 BV510 or CD3 A700 (UCHT1), CD14 BV421 (M5E2), CD19 BV421 (HIB19), CD16 APC-Cy7 (3G8) and CD56 BV605 (NCAM16.2 or B159) to gate on NK cells, and finally fixed, permeabilized (Thermofisher Fix and Perm) and stained with IFN-γ FITC (B27) antibodies to detect intracellular cytokines (all BD Biosciences antibodies). In all assays described above, incubation in the presence of 5 μg/mL of phytohemagglutinin (PHA) or PMA/ionomycin were used as positive controls and unstimulated NK cells or, when applicable, non-pulsed BLCL served as negative controls and for background subtraction. A Fluorescence Minus One (FMO) control and PHA-stimulated PBMCs were used to set the gates for positive cytokine responses. Acquisition of data was performed on a BD LSRII instrument. Data was analyzed using Flow Jo v.10.7.1.

### Generation of primary NK cell clones

Single primary human NK cells were cloned by limiting dilution or single-cell sorting using a protocol adapted from a previously reported method^62^ (Fig. S4). Briefly, NK cells were isolated from PBMC via negative selection (EasySep Cell Separation; StemCell Technologies), added to a mix of irradiated feeders consisting of freshly isolated allogeneic PBMC combined with log-phase-growth RPMI 8866 cells (Sigma-Aldrich) at a 10:1 ratio in cloning medium. Cloning medium is composed of RPMI-1640 supplemented with 5% human serum (Sigma-Aldrich), 1X MEM-NEAA (Gibco), 1X sodium pyruvate (Gibco), 100 μg/mL kanamycin, 300 U/mL Roche recombinant human IL-2 (Sigma-Aldrich) and 1 μg/mL phytohaemagglutinin (PHA; Fisher). The co-cultures were mixed thoroughly and plated at 100 μL/well (1 NK cell/well) in 96-well plates. After 14 days of incubation at 37°C/5% CO2, wells that had outgrowth of cells were transferred to 48-well plates and maintained in NK-cell cloning medium with frequent media exchange (approximately every 3 days). NK cell clones were stained with the following antibodies for flow cytometric phenotyping: BD Biosciences CD3-A700 (UCHT1), CD16-APC-Cy7 (3G8), CD56-BV605 (NCAM16.2), Beckman Coulter NKG2A PE-Cy7 (Z199), and R&D NKG2C-PE (134591). Only NK cell clones that were CD3−CD56+ were used for subsequent assays.

### Human Subjects

De-identified and coded blood samples from the majority of HIV-negative healthy donors used in this study were collected under IRB-approved protocols and delivered to us by Research Blood Components LLC (Watertown, Massachusetts). Additional healthy donors and all PLWH were recruited from outpatient clinics at Massachusetts General Hospital and affiliated Boston-area hospitals or at UAB University Hospital. The respective institutional review boards approved this study, and all subjects gave written informed consent. A total of 4 elite controllers, 27 untreated viremic chronic progressors, and 20 c-ART-treated chronically infected PLWH were studied (Table S1). Elite controllers were defined as having plasma HIV RNA levels of <50 copies/ml in the absence of antiretroviral therapy, on at least three determinations over at least a year of follow-up. Viremic controllers had detectable HIV-1 RNA levels of <2,000 copies/ml. Untreated viremic chronic progressors were defined as subjects having untreated HIV infection for >1 year with plasma viral loads of >2,000 copies/ml for at least 1 year of follow-up; cART treated chronically infected had HIV RNA levels below the limit of detection for the respective available standard assays (e.g., <75 RNA copies/ml by branched DNA assay or <50 copies by PCR). PBMCs were isolated from whole blood by Ficoll-Hypaque density gradient centrifugation, frozen (90% FBS-10% DMSO), and either used immediately or stored in LN2 vapor until analyzed.

### Calcein acetoxymethyl (AM)-based antigen-specific NK cell killing assay for NKCL

Autologous BLCL were stained with 10μm Calcein AM (Invitrogen) for 1h, then washed three times prior to be pulsed with overlapping peptide pools encompassing HIV-1 Consensus B Gag or Env (NIH AIDS Reagent Program), influenza A/California/04/2009(H1N1) NP (PepTivator, Miltenyi Biotec), influenza A/California/08/2009(H1N1) MP1 (PepMix™, JPT Peptide Technologies), CMV pp65 (NIH AIDS Reagent Program), and Myelin-oligodendrocyte glycoprotein (MOG) negative control (PepMix™, JPT Peptide Technologies) or with individual nonameric peptides. Peptide-pulsed BLCL or mock-pulsed controls were then incubated with or without NK cell clones at a 5:1 E:T ratio. Supernatant were harvested after 4h incubation at 37°C/5% CO2. Release of CAM into the supernatant was measured using a fluorescence reader (excitation 485 nm, absorption 530 nm). The percent-specific lysis was calculated as follows: (test release – spontaneous release)/(maximum release - spontaneous release) x 100.

### RNA Isolation

Between 100,000 and 500,000 NKCL were pelleted (500 x g, for 10 minutes) and dry pellets immediately frozen at −80 °C. Pellets were lysed by vortexing for 1 minute in 350uL cold supplemented RLT buffer (RLT + β-MeOH) at 4°C and lysates homogenized using QIAshredder columns (Qiagen). RNA was then extracted from these samples using the RNeasy Plus Micro kit (Qiagen) with on-column DNase digestion. After isolation of total RNA the RNA integrity was analyzed with the RNA 6000 Pico Chip on an Agilent 2100 Bioanalyzer (Agilent Technologies).

### RNA transcriptome analysis

Prior to library generation, RNA was subjected to DNAse I digestion (Thermo Fisher Scientific) followed by RNeasy MinElute column clean up (Qiagen). RNA-Seq libraries were generated using the SMART-Seq v4 Ultra Low Input RNA Kit (Clontech Laboratories) as per the manufacturer’s recommendations. From cDNA final libraries were generated utilizing the Nextera XT DNA Library Preparation Kit (Illumina). Concentrations of the final libraries were measured with a Qubit 2.0 Fluorometer (Thermo Fisher Scientific) and fragment lengths distribution was analyzed with the DNA High Sensitivity Chip on an Agilent 2100 Bioanalyzer (Agilent Technologies). All samples were normalized to 2nM and pooled equimolar. The library pool was sequenced on the NextSeq500 (Illumina) with 1×75bp, with ~14 to 19 mio reads per sample.

Transcript expression was quantified with salmon v. 0.14.0 using default parameters against the GRCh38 human genome reference. Differential gene expression (DGE) was performed using DESeq2 v. 1.28.1 with a previously described protocol^63,64^. An adjusted p-value cutoff of 0.05 was used to determine significance. Differentially regulated genes were selected by calculation of a false-discovery rate below 0.05 and log2 fold change values greater than or equal to 1.5 and less than or equal to −1.5. Gene symbols and Entrez identifiers were mapped to Ensembl gene identifiers using biomaRt v. 2.44.1d^65,66^. Gene set enrichment analysis (GSEA) was used with Entrez IDs and ranked based on the following formula:

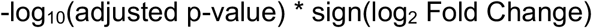

Msigdbr v. 7.1.1 was used to retrieve Gene Ontology (GO) and Hallmark pathways^67^ and GSEA was performed using the ranked gene list and pathways with the function fgseaSimple() from the fgsea v. 1.14.0 package with default parameters^68^. Heatmaps were generated using pheatmap v. 1.0.12^69^. Volcano plots and GSEA results were generated using ggplot2 v. 3.3.2^70^.

### Advanced Polychromatic flow cytometry for the phenotypic characterization of NKCL

NKCL previously frozen in liquid nitrogen were stained with panel of antibodies detailed in Table S2. Data were acquired on a FACS Symphony A5 flow cytometer (BD Biosciences) equipped with five lasers (UV, 355 nm; violet, 405 nm; blue, 488; yellow/green, 561 nm; red, 637 nm; all tuned at 100 mW, except UV tuned at 60 mW). Flow Cytometry Standard (FCS) files were exported as FCS3.0 from DiVa 8.0.1. FCS files were loaded and compensated in FlowJo by using single-stained controls (Compbeads incubated with fluorescently conjugated antibodies). After compensation and gating, live NKCL were exported as FCS3.0 files for subsequent t-SNE analysis using CytoDRAV (https://github.com/ReevesLab/CytoDRAV).

### HLA-E stabilization assay

A selection of nonameric peptides derived from subtype B HIV Gag (60) (GenBank accession number ABR15476.1) and Env (84) (UniProtKB accession number A0A160I6K4) as well as influenza A/California/04/2009(H1N1) NP (23) (UniProtKB accession number C3W5S2) predicted to have strong and weak binding to HLA-E*01:01 by the NetMHCpan 4.0 and Immune Epitope Database servers were synthesized using the PEPotec Immuno Custom Peptide Libraries service provided by Thermofisher Scientific (Table S3). Analyses were conducted using influenza NP, HIV Gag and HIV Env sequences matching those used to generate the peptide pools used in our studies. The ability of individual peptides to stabilize surface expression of human HLA-E was assessed using K562-HLA-E 01:01 transfectants cultured at 26°C for 26h prior to be pulsed with 10μg/mL of individual peptides for 16h at 26°C. Cell surface expression levels of HLA-E were assessed by flow cytometry staining using anti-HLA-E-BV421 antibody (3D12). HLA-E expression was compared to that upon pulsing with positive control peptide VL9 (VMAPRTLIL), a canonical CMV UL40/HLA-Cw3 leader sequence-derived peptide that stabilizes HLA-E, and negative control peptide NLVPMVATV, a CMV pp65-derived nonamer that does not stabilize HLA-E. Data were acquired on an LSRII instrument (BD Biosciences), and analyzed using FlowJo software v10.7.1 (Treestar).

## Acknowledgements

We thank all the patients who contributed to this study making this work possible.

## Author Contributions

S.J. and R.K.R. designed the study, analyzed data and generated the final figures. O.L., T.Y., S.S., S.S., R.J., G.T., A.W., P.J.T., H.L.D. and J.G. performed the experiments and analyzed data. A.G. performed the RNAseq with input from M.A. K.K. performed the bioinformatic analyses. P.G. and M.A. provided access to human samples. M.M.T. provided crucial reagents. S.J. and R.K.R. wrote the manuscript with contributions of all authors.

---

**Supplementary Table 1.**
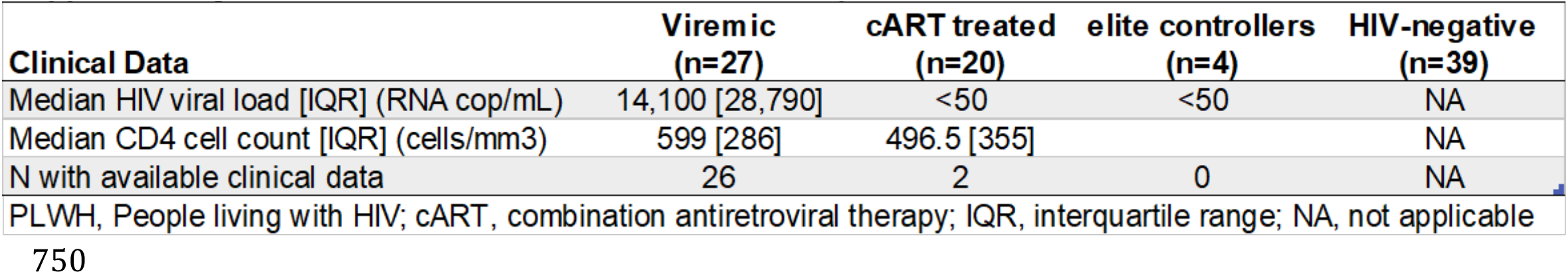
Characteristics of PL WI-I and healthy donors

**Supplementary Table 2.** 28-color panel for NKCL phenotyping

| **Target** | **Fluorochrome** | **Clone** | **Manufacturer** |
| --- | --- | --- | --- |
| LIVE cells | Blue Live/Dead |  | Invitrogen |
| CD19 | PE-CY5.5 | J3-119 | BECKMAN COULTER |
| CD14 | BUV615 | M5E2 | BD Pharmingen |
| CD3 | BUV496 | UCHT1 | BD Pharmingen |
| CD4 | BB700 | L200 | BD Pharmingen |
| CD56 | BUV737 | NCAM16.2 | BD Pharmingen |
| CD16 | BUV563 | 3G8 | BD Pharmingen |
| NKP46 | BV711 | 9E2/NKp46 | BD Pharmingen |
| NKG2D | BB790 | 1D11 | BD Pharmingen |
| NKG2C | Biotin + SA BUV395 | REA205 | MILTENYI |
| NKG2A | PE-CY7 | Z199 | BECKMAN COULTER |
| KIR3DL1/S1 | VioBlue | REA168 | MILTENYI |
| KIR2DL1/S1/S3/S5 | FITC | HP-MA4 | BIOLEGEND |
| KIR2DS4 | APC-Vio770 | REA284 | MILTENYI |
| CD85J | PE | GHI/75 | BD Pharmingen |
| $\alpha 4\beta 7$ | APC | | NHP Reagent Resource |
| CXCR6 | BV786 | 13B 1E5 | BD Pharmingen |
| CCR5 | PE-CY5 | 2D7/CCR5 | BD Pharmingen |
| CCR7 | APC-R700 | 3D12 | BD Pharmingen |
| 2B4 | BV650 | 2-69 | BD Pharmingen |
| CD2 | BV510 | 2H7 | BD Pharmingen |
| CD8 | BV570 | RPA-T8 | BIOLEGEND |
| CD57 | BB630 | NK-1 | BD Pharmingen |
| KLRG1 | PE-Dazzle594 | 2F1/KLRG1 | BIOLEGEND |
| PD1 | BV605 | EH12.1 | BD Pharmingen |
| TIM-3 | BV750 | 7D3 | BD Pharmingen |
| HLA-DR | BUV661 | G46-6 | BD Pharmingen |
| IL-7R | BUV805 | HIL-7R-M21 | BD Pharmingen |

**Supplementary Table 3.**
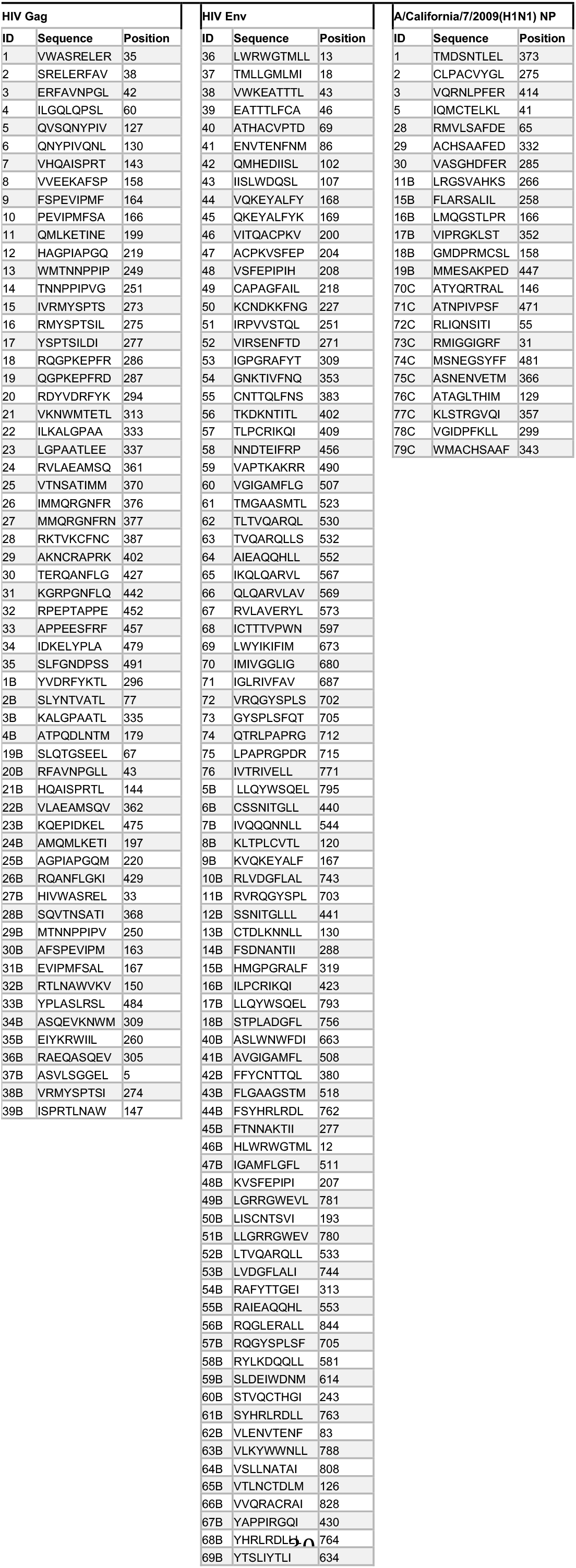
HIV- and influenza-derived nonamers tested for binding to HLA-E*01:01

## SUPPLEMENTARY FIGURE LEGENDS

**Supplementary Fig. 1.**
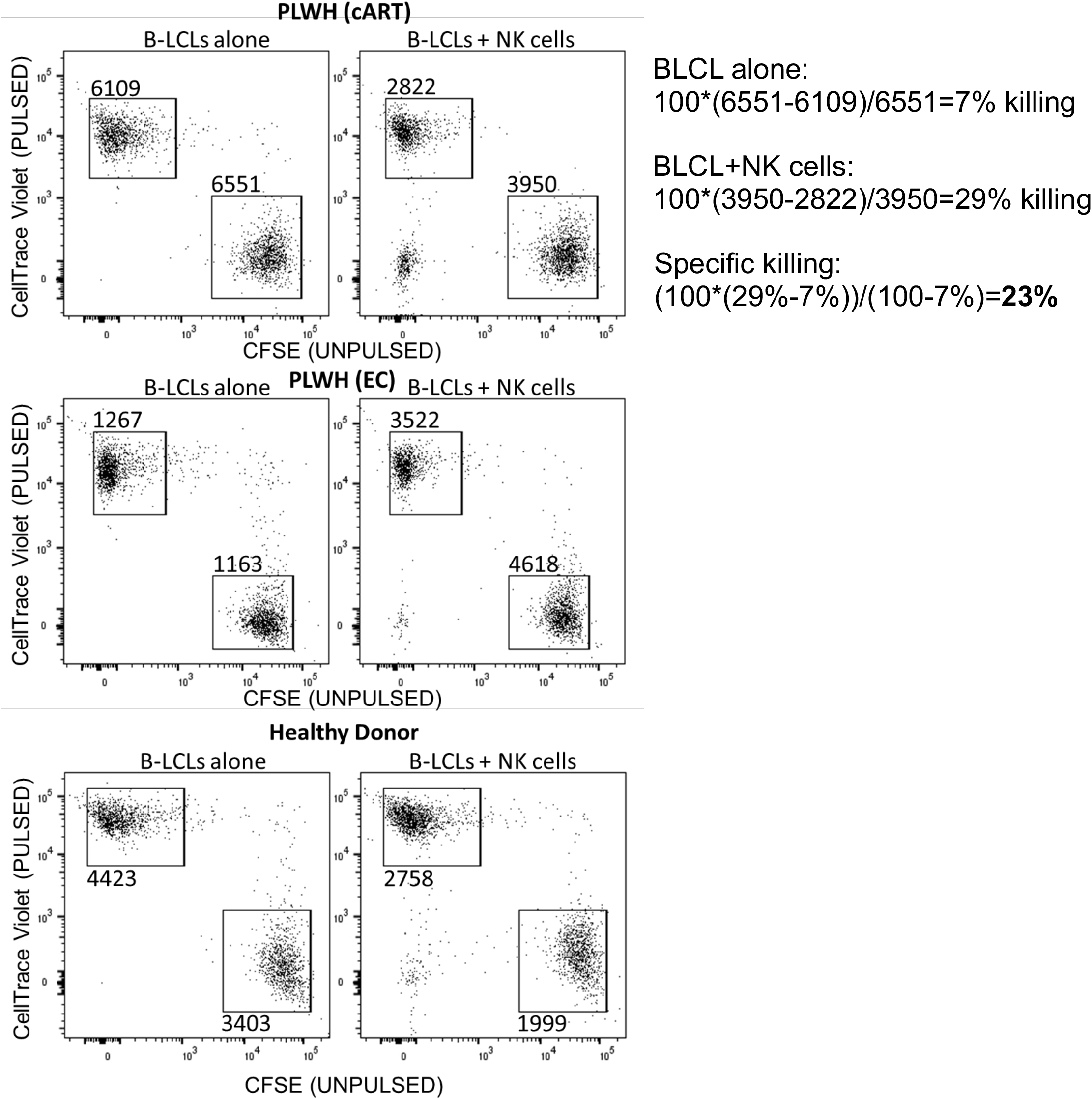
Representative flow cytometry plots depicting CellTrace Violet-stained HIV Gag peptide-pulsed and CFSE-stained unpulsed BLCL from a cART-treated PLWH (upper panels), an elite controller (EC) (middle panels) and a healthy donor (lower panels) alone (left panels) or in the presence of autologous purified NK cells (right panels), after gating on CD3neg CD19pos lymphocytes. Representative example of specific lysis calculation is provided based on the following formula: (% sample lysis with NK effectors - % basal lysis without NK effectors) / (100 - % basal lysis without NK effectors).

**Supplementary Fig. 2.**
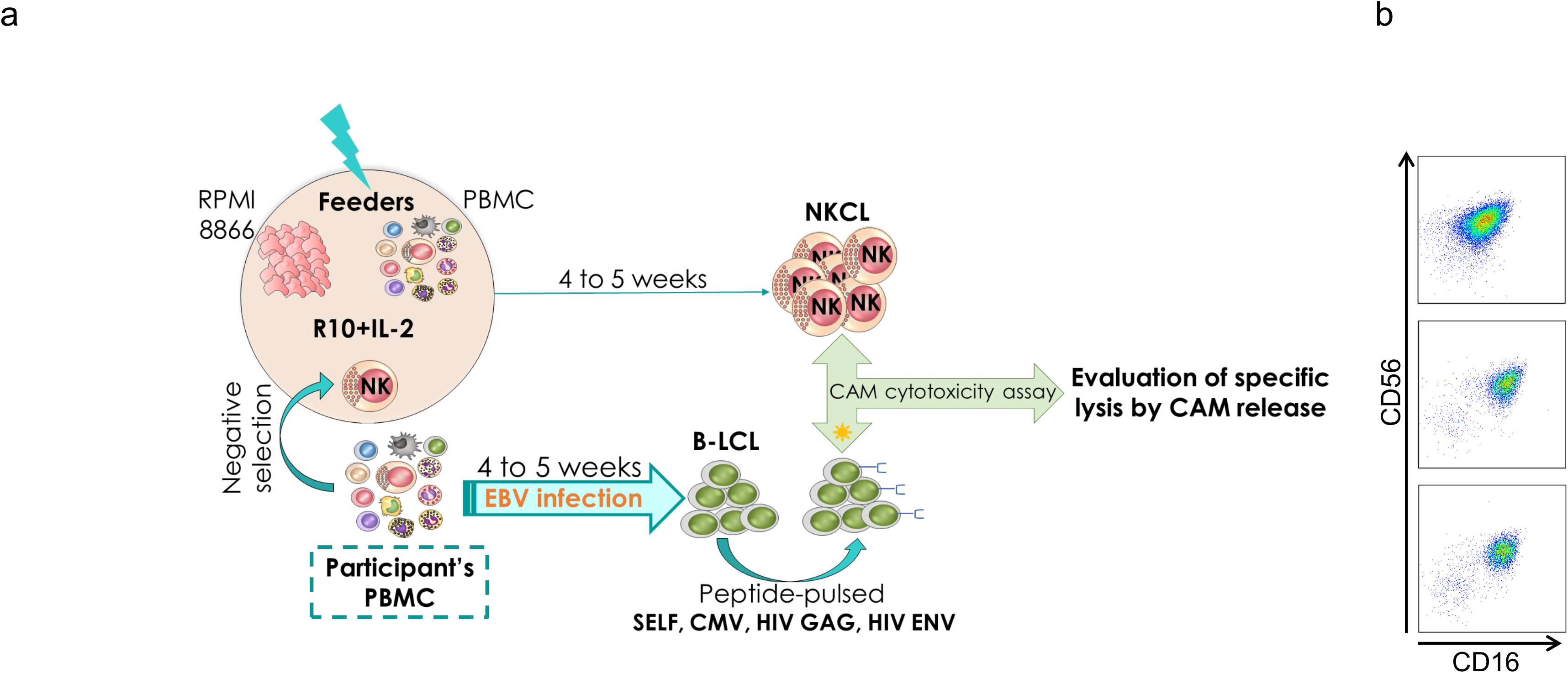
**a,** Graphical design of the methodology to generate NKCL and evaluate their cytotoxic potential against autologous BLCL presenting viral antigens. **b**, Representative flow cytometry plots showing CD16 and CD56 expression of three different NKCL generated from PBMC of one single individual.

**Supplementary Fig. 3.**
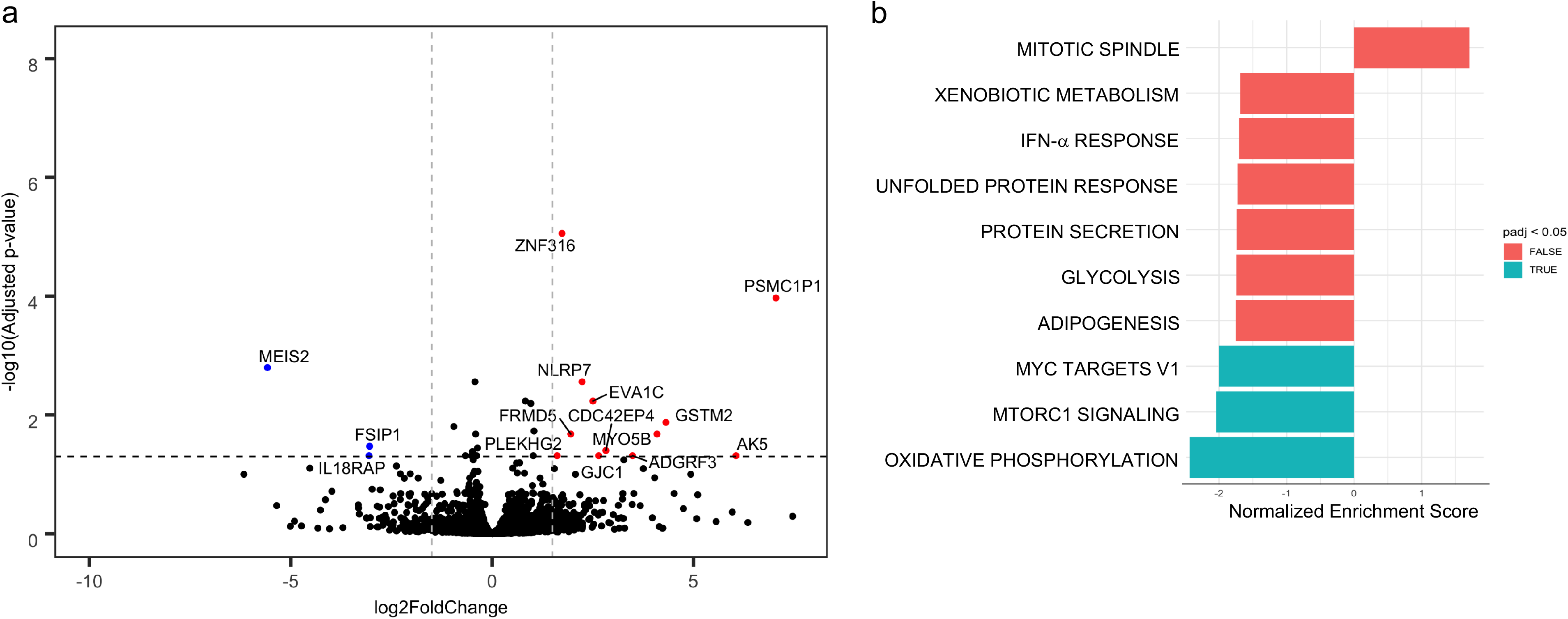
**a,** Volcano plot depicting 15 differentially expressed genes distinguishing HIV-specific NKCL (n=8) and NKCL not reacting to HIV Gag/Env (n=7) from the same untreated viremic PLWH. Red and blue dots represent significantly upregulated and downregulated genes, respectively. Differentially regulated genes were selected by calculation of a false-discovery rate below 0.05 and log2 fold change values greater than or equal to 1.5 and less than or equal to −1.5. **b,** Selected top 10 significant Hallmark pathways for differentially expressed genes in HIV-specific NKCL compared to NKCL not reacting to HIV Gag/Env, as evaluated by Gene Set Enrichment Analysis software using the fgsea v. 1.14.0 package with pathways selected using Msigdbr v. 7.1.1 package. Blue shading indicates an adjusted p-value less than the 0.05 cutoff. Red shading indicates an adjusted p-value greater than 0.05.

**Supplementary Fig. 4.**
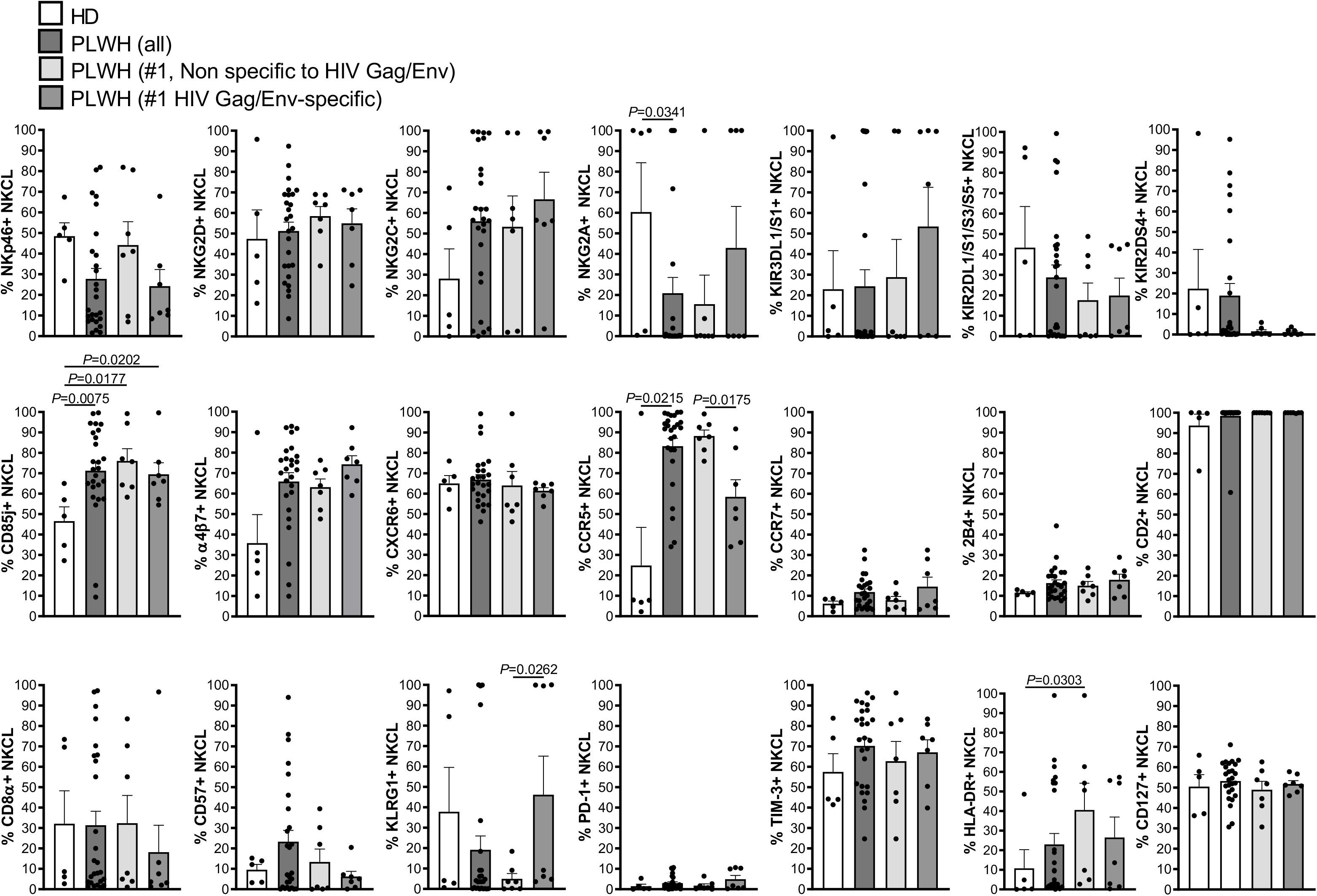
Bar graphs show mean + SEM proportions of NKCL expressing indicated surface markers evaluated by flow cytometry and comparing 5 NKCL from 2 healthy donors (HD, white bars) with 14 NKCL from one untreated viremic PLWH (darker grey bars), including 7 HIV Gag/Env-specific NKCL (dark grey bars) and 7 NKCL that did not react to HIV Gag/Env (light grey bars).

**Supplementary Fig. 5.**
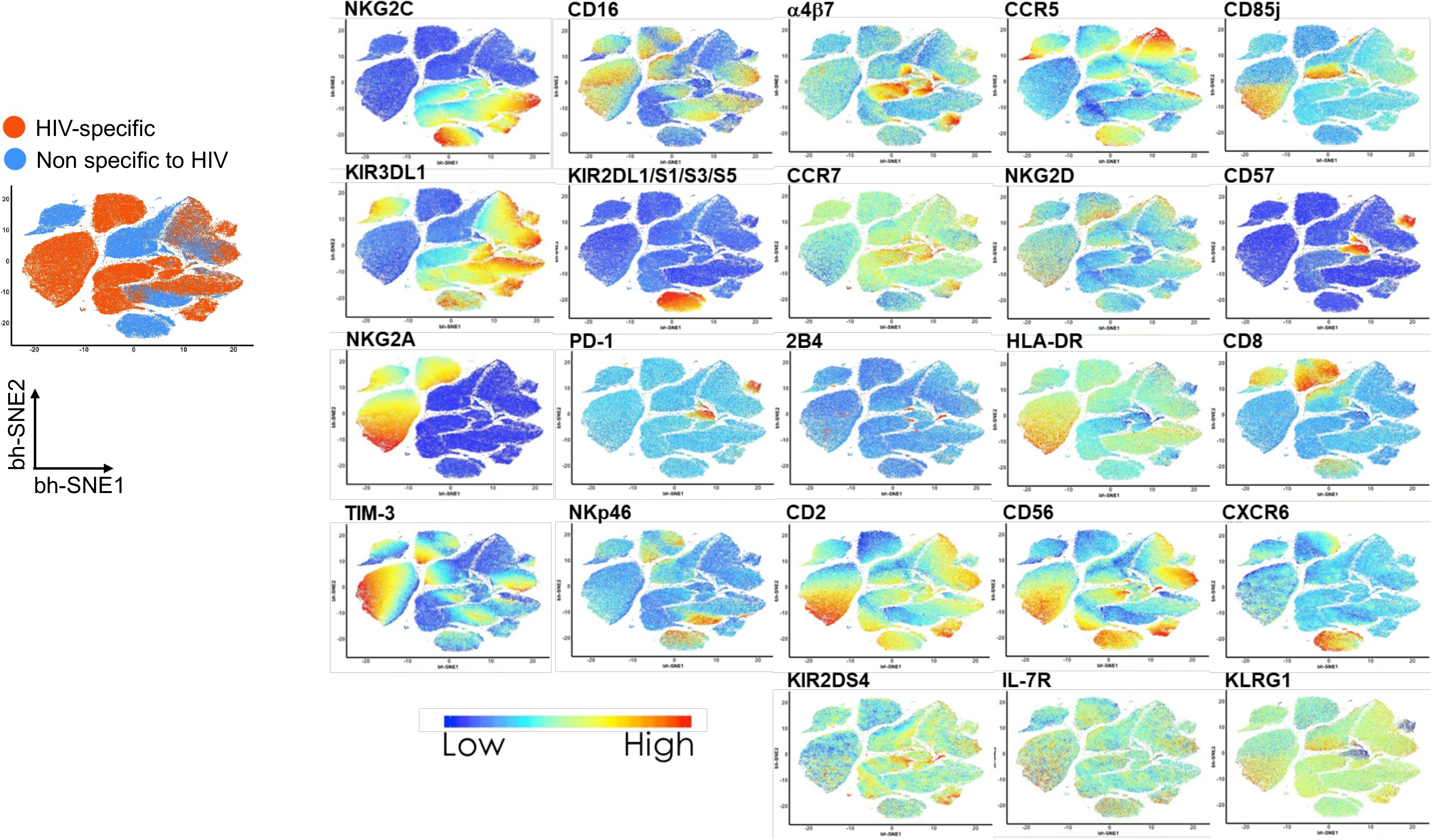
Multidimensional data analysis of 14 NKCL from one untreated viremic PLWH. Live NKCL were analyzed by *t*-SNE with bh-SNE to generate plots clustering cells with similar expression profiles. Relative expression of indicated NK cell markers visualized over the bh-SNE plots is displayed as a colorimetric scale of blue (low expression) to red (high expression).

**Supplementary Fig. 6.**
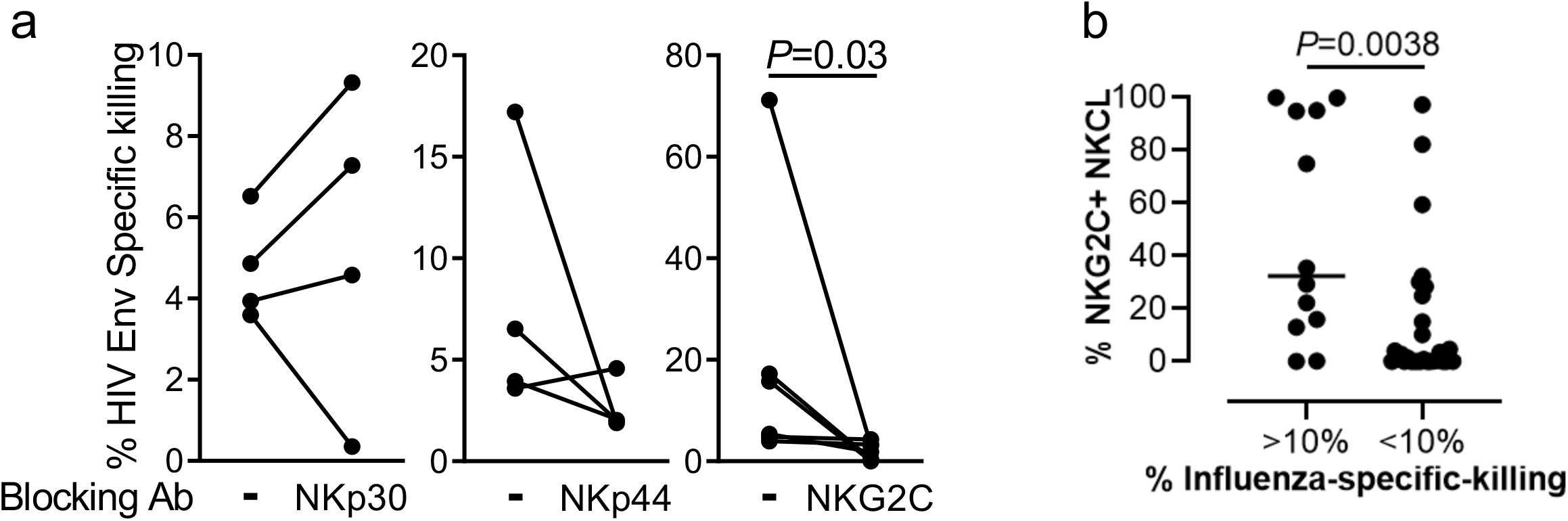
**a,** Antigen-specific killing of HIV Env-pulsed BLCL by NKCL in the presence of control or isotype control, NKp30, NKp44 or NKG2C blocking antibodies. **b**, Percentages of NKG2C+ NKCL, comparing NKCL with more (>10%) or less (<10%) than 10% influenza-specific killing for 41 NKCL from 9 healthy donors. For each NKCL, the highest antigen-specific killing against influenza HA, NP and MP1 was considered. Bar indicates the median.

**Supplementary Fig. 7.**
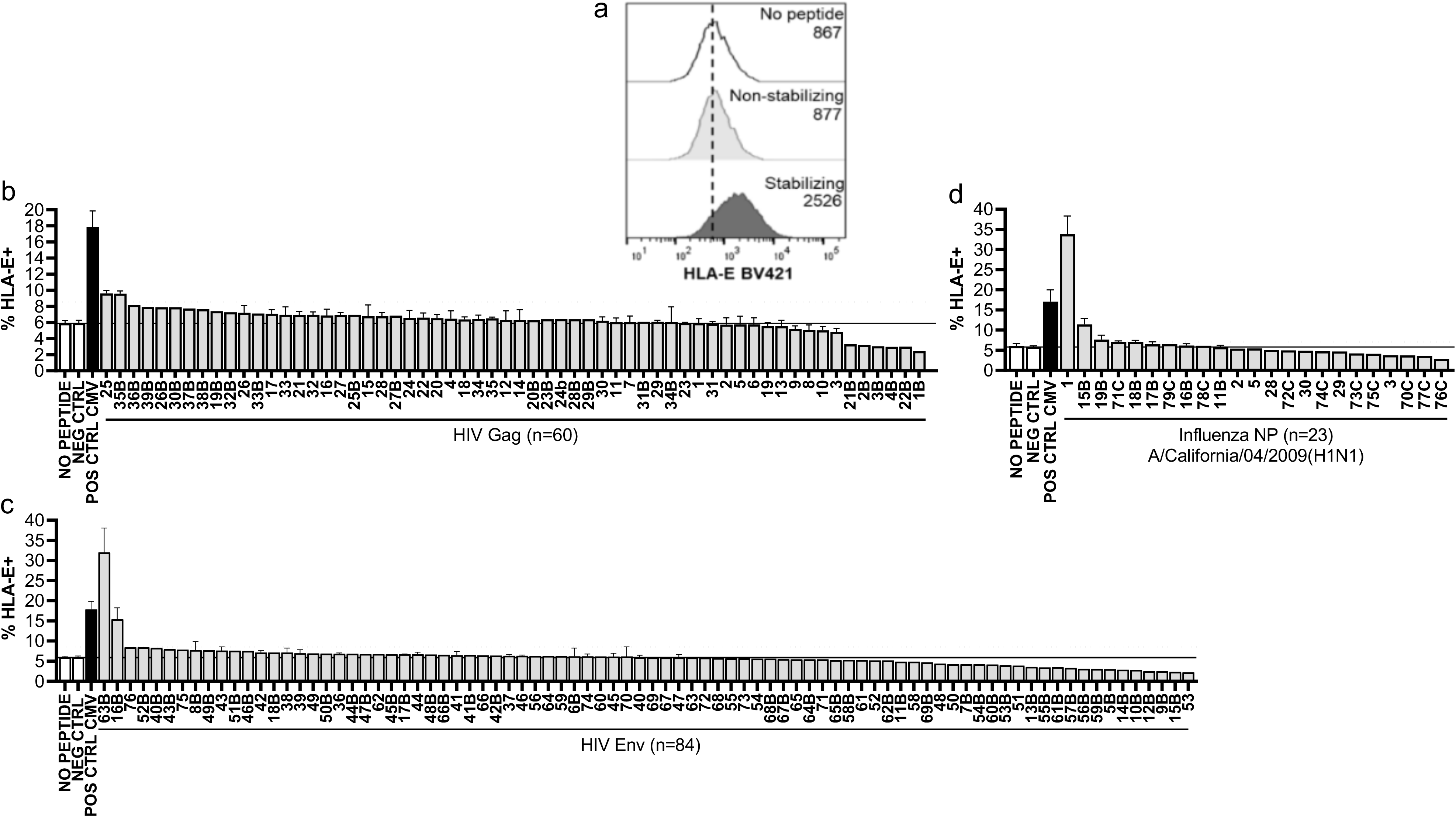
**a**, Representative flow cytometry plots showing HLA-E expression on K562 stably expressing HLA-E*01:01 cells that were either left unpulsed (upper panel) or pulsed for 16 h at 26°C with 5ug/mL of CMV-derived NLVPMVATV that do not stabilize HLA-E (middle panel), or with VMAPRTLIL, a CMV/HLA-Cw3 leader sequence-derived peptide that stabilizes HLA-E (lower panel). Bar graphs represent mean percentages +SEM of HLA-E+ K562 cells pooled from at least 2 distinct experiments after pulsing with NLVPMVATV (NEG CTRL), VMAPRTLIL (POS CTRL CMV), 60 single nonameric peptides derived from HIV Gag (**b**), 84 single nonameric peptides derived from HIV Env (**c**), or 23 single peptides derived from NP A/California/04/2009(pH1N1) (**d**) with strong binding prediction scores based on the NetMHC pan 4.0 and IEDB epitope prediction tools. HLA-E surface stabilization was assessed using anti-HLA-E antibody (3D12). Dead cells were excluded. The dotted line marks the cut off for increased HLA-E expression, defined as the mean value plus 2 standard deviations of non-stabilizing peptides.

## References

1 Florez-Alvarez, L., Hernandez, J. C. & Zapata, W. NK Cells in HIV-1 Infection: From Basic Science to Vaccine Strategies. Front Immunol 9, 2290, doi:10.3389/fimmu.2018.02290 (2018).

2 Scully, E. & Alter, G. NK Cells in HIV Disease. Current HIV/AIDS reports 13, 85–94, doi:10.1007/s11904-016-0310-3 (2016).

3 Jost, S. & Altfeld, M. Control of Human Viral Infections by Natural Killer Cells. Annual review of immunology, doi:10.1146/annurev-immunol-032712-100001 (2013).

4 Shah, S. V. et al. CMV Primes Functional Alternative Signaling in Adaptive Deltag NK Cells but Is Subverted by Lentivirus Infection in Rhesus Macaques. Cell Rep 25, 2766–2774 e2763, doi:1016/j.celrep.2018.11.020 (2018).

5 O’Leary, J. G., Goodarzi, M., Drayton, D. L. & von Andrian, U. H. T cell- and B cell-independent adaptive immunity mediated by natural killer cells. Nat Immunol 7, 507–516 (2006).

6 Sun, J. C., Beilke, J. N. & Lanier, L. L. Adaptive immune features of natural killer cells. Nature 457, 557–561, doi:10.1038/nature07665 (2009).

7 Paust, S. et al. Critical role for the chemokine receptor CXCR6 in NK cell-mediated antigen-specific memory of haptens and viruses. Nat Immunol 11, 1127–1135, doi:10.1038/ni.1953 (2010).

8 Gillard, G. O. et al. Thy1+ NK [corrected] cells from vaccinia virus-primed mice confer protection against vaccinia virus challenge in the absence of adaptive lymphocytes. PLoS pathogens 7, e1002141, doi:10.1371/journal.ppat.1002141 (2011).

9 Foley, B. et al. Cytomegalovirus reactivation after allogeneic transplantation promotes a lasting increase in educated NKG2C+ natural killer cells with potent function. Blood 119, 2665–2674, doi:10.1182/blood-2011-10-386995 (2012).

10 Peng, H. et al. Liver-resident NK cells confer adaptive immunity in skin-contact inflammation. The Journal of clinical investigation 123, 1444–1456, doi:10.1172/JCI66381 (2013).

11 Lee, J. et al. Epigenetic modification and antibody-dependent expansion of memory-like NK cells in human cytomegalovirus-infected individuals. Immunity 42, 431–442, doi:10.1016/j.immuni.2015.02.013 (2015).

12 Paust, S., Blish, C. A. & Reeves, R. K. Redefining Memory: Building the Case for Adaptive NK Cells. Journal of virology 91, doi:10.1128/JVI.00169-17 (2017).

13 Reeves, R. K. et al. Antigen-specific NK cell memory in rhesus macaques. Nature immunology 16, 927–932, doi:10.1038/ni.3227 (2015).

14 Li, T. et al. Respiratory Influenza Virus Infection Induces Memory-like Liver NK Cells in Mice. Journal of immunology 198, 1242–1252, doi:10.4049/jimmunol.1502186 (2017).

15 Hammer, Q. et al. Peptide-specific recognition of human cytomegalovirus strains controls adaptive natural killer cells. Nat Immunol 19, 453–463, doi:10.1038/s41590-018-0082-6 (2018).

16 Gardiner, C. M. & Mills, K. H. The cells that mediate innate immune memory and their functional significance in inflammatory and infectious diseases. Semin Immunol 28, 343–350, doi:10.1016/j.smim.2016.03.001 (2016).

17 Blok, B. A., Arts, R. J., van Crevel, R., Benn, C. S. & Netea, M. G. Trained innate immunity as underlying mechanism for the long-term, nonspecific effects of vaccines. J Leukoc Biol 98, 347–356, doi:10.1189/jlb.5RI0315-096R (2015).

18 Nikzad, R. et al. Human natural killer cells mediate adaptive immunity to viral antigens. Sci Immunol 4, doi:10.1126/sciimmunol.aat8116 (2019).

19 Wijaya, R. S. et al. HBV vaccination and HBV infection induces HBV-specific natural killer cell memory. Gut, doi:gutjnl-2019-319252 (2020).

20 Stary, V. et al. A discrete subset of epigenetically primed human NK cells mediates antigen-specific immune responses. Sci Immunol 5, doi:10.1126/sciimmunol.aba6232 (2020).

21 Ma, M. et al. NKG2C(+)NKG2A(-) Natural Killer Cells are Associated with a Lower Viral Set Point and may Predict Disease Progression in Individuals with Primary HIV Infection. Front Immunol 8, 1176, doi:10.3389/fimmu.2017.01176 (2017).

22 Gondois-Rey, F. et al. NKG2C(+) memory-like NK cells contribute to the control of HIV viremia during primary infection: Optiprim-ANRS 147. Clin Transl Immunology 6, e150, doi:10.1038/cti.2017.22 (2017).

23 Thomas, R. et al. NKG2C deletion is a risk factor of HIV infection. AIDS Res Hum Retroviruses 28, 844–851, doi:10.1089/AID.2011.0253 (2012).

24 Peppa, D. et al. Adaptive Reconfiguration of Natural Killer Cells in HIV-1 Infection. Front Immunol 9, 474, doi:10.3389/fimmu.2018.00474 (2018).

25 Suliman, S. et al. Bacillus Calmette-Guerin (BCG) Revaccination of Adults with Latent Mycobacterium tuberculosis Infection Induces Long-Lived BCG-Reactive NK Cell Responses. Journal of immunology 197, 1100–1110, doi:10.4049/jimmunol.1501996 (2016).

26 Rolle, A., Meyer, M., Calderazzo, S., Jager, D. & Momburg, F. Distinct HLA-E Peptide Complexes Modify Antibody-Driven Effector Functions of Adaptive NK Cells. Cell Rep 24, 1967–1976 e1964, doi:10.1016/j.celrep.2018.07.069 (2018).

27 Pereyra, F. et al. Genetic and immunologic heterogeneity among persons who control HIV infection in the absence of therapy. The Journal of infectious diseases 197, 563–571, doi:10.1086/526786 (2008).

28 Guma, M. et al. Imprint of human cytomegalovirus infection on the NK cell receptor repertoire. Blood 104, 3664–3671 (2004).

29 Hendricks, D. W. et al. Cutting edge: NKG2C(hi)CD57+ NK cells respond specifically to acute infection with cytomegalovirus and not Epstein-Barr virus. Journal of immunology 192, 4492–4496, doi:10.4049/jimmunol.1303211 (2014).

30 Lopez-Verges, S. et al. Expansion of a unique CD57(+)NKG2Chi natural killer cell subset during acute human cytomegalovirus infection. Proceedings of the National Academy of Sciences of the United States of America 108, 14725–14732, doi:10.1073/pnas.1110900108 (2011).

31 Schlums, H. et al. Cytomegalovirus infection drives adaptive epigenetic diversification of NK cells with altered signaling and effector function. Immunity 42, 443–456, doi:10.1016/j.immuni.2015.02.008 (2015).

32 Tomasec, P. et al. Surface expression of HLA-E, an inhibitor of natural killer cells, enhanced by human cytomegalovirus gpUL40. Science 287, 1031, doi:science.287.5455.1031 (2000).

33 Pietra, G. et al. HLA-E-restricted recognition of cytomegalovirus-derived peptides by human CD8+ cytolytic T lymphocytes. Proc Natl Acad Sci U S A 100, 10896–10901, doi:pnas.1834449100 (2003).

34 Nattermann, J. et al. The HLA-A2 restricted T cell epitope HCV core 35-44 stabilizes HLA-E expression and inhibits cytolysis mediated by natural killer cells. Am J Pathol 166, 443–453, doi:10.1016/S0002-9440(10)62267-5 (2005).

35 Garcia, P. et al. Human T cell receptor-mediated recognition of HLA-E. Eur J Immunol 32, 936–944, doi:10.1002/1521-4141(200204)32:4<936::AID-IMMU936>3.0.CO;2-M (2002).

36 Nattermann, J. et al. HIV-1 infection leads to increased HLA-E expression resulting in impaired function of natural killer cells. Antivir Ther 10, 95–107 (2005).

37 Davis, Z. B. et al. A Conserved HIV-1-Derived Peptide Presented by HLA-E Renders Infected T-cells Highly Susceptible to Attack by NKG2A/CD94-Bearing Natural Killer Cells. PLoS pathogens 12, e1005421, doi:10.1371/journal.ppat.1005421 (2016).

38 Hannoun, Z. et al. Identification of novel HIV-1-derived HLA-E-binding peptides. Immunol Lett 202, 65–72, doi:10.1016/j.imlet.2018.08.005 (2018).

39 Aktas, E., Erten, G., Kucuksezer, U. C. & Deniz, G. Natural killer cells: versatile roles in autoimmune and infectious diseases. Expert Rev Clin Immunol 5, 405–420, doi:10.1586/eci.09.27 (2009).

40 Alter, G., Malenfant, J. M. & Altfeld, M. CD107a as a functional marker for the identification of natural killer cell activity. J Immunol Methods 294, 15–22, doi:10.1016/j.jim.2004.08.008 (2004).

41 Hansen, S. G. et al. Profound early control of highly pathogenic SIV by an effector memory T-cell vaccine. Nature 473, 523–527, doi:10.1038/nature10003 (2011).

42 Hansen, S. G. et al. A live-attenuated RhCMV/SIV vaccine shows long-term efficacy against heterologous SIV challenge. Sci Transl Med 11, doi:10.1126/scitranslmed.aaw2607 (2019).

43 Wu, H. L. et al. The Role of MHC-E in T Cell Immunity Is Conserved among Humans, Rhesus Macaques, and Cynomolgus Macaques. J Immunol 200, 49–60, doi:10.4049/jimmunol.1700841 (2018).

44 Paust, S., Blish, C. A. & Reeves, R. K. Redefining Memory: Building the Case for Adaptive NK Cells. J Virol 91, doi:10.1128/JVI.00169-17 (2017).

45 Fogel, L. A., Sun, M. M., Geurs, T. L., Carayannopoulos, L. N. & French, A. R. Markers of nonselective and specific NK cell activation. J Immunol 190, 6269–6276, doi:10.4049/jimmunol.1202533 (2013).

46 Venkatasubramanian, S. et al. IL-21-dependent expansion of memory-like NK cells enhances protective immune responses against Mycobacterium tuberculosis. Mucosal Immunol 10, 1031–1042, doi:10.1038/mi.2016.105 (2017).

47 Reeves, R. K., Evans, T. I., Gillis, J. & Johnson, R. P. Simian immunodeficiency virus infection induces expansion of alpha4beta7+ and cytotoxic CD56+ NK cells. Journal of virology 84, 8959–8963, doi:10.1128/JVI.01126-10 (2010).

48 Sips, M. et al. Altered distribution of mucosal NK cells during HIV infection. Mucosal immunology 5, 30–40, doi:10.1038/mi.2011.40 (2012).

49 Khakoo, S. I. et al. HLA and NK cell inhibitory receptor genes in resolving hepatitis C virus infection. Science 305, 872–874, doi:10.1126/science.1097670 (2004).

50 Malnati, M. S. et al. Activating Killer Immunoglobulin Receptors and HLA-C: a successful combination providing HIV-1 control. Sci Rep 7, 42470, doi:10.1038/srep42470 (2017).

51 Martin, M. P. et al. Epistatic interaction between KIR3DS1 and HLA-B delays the progression to AIDS. Nat Genet 31, 429–434 (2002).

52 Martin, M. P. et al. Innate partnership of HLA-B and KIR3DL1 subtypes against HIV-1. Nature genetics 39, 733–740, doi:10.1038/ng2035 (2007).

53 Fadda, L. et al. Common HIV-1 peptide variants mediate differential binding of KIR3DL1 to HLA-Bw4 molecules. Journal of virology 85, 5970–5974, doi:10.1128/JVI.00412-11 (2011).

54 Peruzzi, M., Parker, K. C., Long, E. O. & Malnati, M. S. Peptide sequence requirements for the recognition of HLA-B*2705 by specific natural killer cells. J Immunol 157, 3350–3356 (1996).

55 Stewart-Jones, G. B. et al. Crystal structures and KIR3DL1 recognition of three immunodominant viral peptides complexed to HLA-B*2705. Eur J Immunol 35, 341–351, doi:10.1002/eji.200425724 (2005).

56 Saunders, P. M. et al. The interaction of KIR3DL1*001 with HLA class I molecules is dependent upon molecular microarchitecture within the Bw4 epitope. J Immunol 194, 781–789, doi:10.4049/jimmunol.1402542 (2015).

57 O’Connor, G. M. et al. Mutational and structural analysis of KIR3DL1 reveals a lineage-defining allotypic dimorphism that impacts both HLA and peptide sensitivity. J Immunol 192, 2875–2884, doi:10.4049/jimmunol.1303142 (2014).

58 Jiang, C. et al. Distinct viral reservoirs in individuals with spontaneous control of HIV-1. Nature 585, 261–267, doi:10.1038/s41586-020-2651-8 (2020).

59 Migueles, S. A. & Connors, M. Success and failure of the cellular immune response against HIV-1. Nat Immunol 16, 563–570, doi:10.1038/ni.3161 (2015).

60 Gaiha, G. D. et al. Structural topology defines protective CD8(+) T cell epitopes in the HIV proteome. Science 364, 480–484, doi:10.1126/science.aav5095 (2019).

61 Migueles, S. A. et al. Lytic granule loading of CD8+ T cells is required for HIV-infected cell elimination associated with immune control. Immunity 29, 1009–1021, doi:10.1016/j.immuni.2008.10.010 (2008).

62 Cella, M. & Colonna, M. Cloning human natural killer cells. Methods in molecular biology 121, 1–4, doi:10.1385/1-59259-044-6:1 (2000).

63 Love, M. I., Huber, W. & Anders, S. Moderated estimation of fold change and dispersion for RNA-seq data with DESeq2. Genome Biol 15, 550, doi:10.1186/s13059-014-0550-8 (2014).

64 Love, M. I., Huber, W. & Anders, S. Analyzing RNA-seq data with DESeq2. Retrieved from https://bioconductor.org/packages/release/bioc/vignettes/DESeq2/inst/doc/DESeq2.html. (2020).

65 Durinck, S. et al. BioMart and Bioconductor: a powerful link between biological databases and microarray data analysis. Bioinformatics 21, 3439–3440, doi:10.1093/bioinformatics/bti525 (2005).

66 Durinck, S., Spellman, P. T., Birney, E. & Huber, W. Mapping identifiers for the integration of genomic datasets with the R/Bioconductor package biomaRt. Nat Protoc 4, 1184–1191, doi:10.1038/nprot.2009.97 (2009).

67 Dolgalev, I. msigdbr: MSigDB Gene Sets for Multiple Organisms in a Tidy Data Format (Version 7.1.1). Retrieved from https://CRAN.R-project.org/package=msigdbr. (2020).

68 Korotkevich, G., Sukhov, V. & Sergushichev, A. Fast gene set enrichment analysis. BioRxiv. Preprint published online October 22, 2019, doi:https://doi.org/10.1101/060012 (2019).

69 Kolde, R. pheatmap: Pretty Heatmaps (Version 1.0.12). Retrieved from https://CRAN.R-project.org/package=pheatmap. (2019).

70 Wickham, H. ggplot2: Elegant graphics for data analysis (Second edition). Springer. (2016).

